# Changes in neural progenitor lineage constituent during astrocytic differentiation of human iPSCs

**DOI:** 10.1101/2024.01.22.575551

**Authors:** Zongze Li, Lucia Fernandez Cardo, Michal Rokicki, Jimena Monzón-Sandoval, Viola Volpato, Frank Wessely, Caleb Webber, Meng Li

## Abstract

The regional specificity of stem cell-derived astrocytes is believed to be an important prerequisite for their application in disease modelling and cell-based therapies. Due to the lack of subtype-defining markers for astrocytes in different regions of the brain, the regional identity of *in vitro*-derived astrocytes is often declared by the dominant positional characteristics of their antecedent neural progenitors, patterned to a fate of interest, with the assumption that the positional trait is preserved by the derived astrocytes via linear descent. Using a human induced pluripotent stem cell line designed for tracing derivatives of LMX1A^+^ cells combined with a ventral midbrain induction paradigm, we show that astrocytes originating from LMX1A^+^ progenitors can only be generated if these progenitors are purified prior to the astrocyte differentiation process, or their progenies are gradually lost to progenies of LMX1A^-^ progenitors. This finding indicates that the lineage composition of iPSC-derived astrocytes may not accurately recapitulate the founder progenitor population. Using deep single-cell RNA sequencing, we identified distinct transcriptomic signatures in astrocytes derived from the LMX1A^+^ progenitor cells. Our study highlights the need for rigorous characterization of pluripotent stem cell-derived regional astrocytes, and provides a resource for assessing LMX1A^+^ ventral midbrain progenitor-derived human astrocytes.

## Introduction

Astrocytes are the most abundant cell type in the brain. They play important roles in the central nervous system in supporting neuronal survival and synaptic activities, including regulation of ionic homeostasis, providing energetic support, elimination of oxidative stress, and neurotransmitter removal and recycling (Verkhratsky and Nedergaard 2018). Abnormalities in astrocytes have been linked to various neurodegenerative and neurodevelopmental disorders such as Parkinson’s disease, Alzheimer’s disease, Huntington’s disease, autism spectrum disorders, and Alexander’s disease (Molofsky et al. 2012; Phatnani and Maniatis 2015; Booth et al. 2017). Therefore, there is a growing interest in using human pluripotent stem cell (PSC)-derived astrocytes for in vitro disease modeling (Chandrasekaran et al. 2016).

Contrary to the widely held belief that astrocytes in the brain are largely identical, recent studies have revealed a diversity in their transcriptomic profiles, physiological properties, and functions (Oberheim et al. 2009; Schober et al. 2022). Single-cell and spatial transcriptomic studies have identified several astrocyte subpopulations in the mouse cortex (Zhu et al. 2018; Batiuk et al. 2020; Bayraktar et al. 2020). In humans, although astrocyte heterogeneity remains largely elusive, heterogeneity in radial glia across brain regions and within the midbrain has been reported (La Manno et al. 2016; Li et al. 2023). Furthermore, different molecular and physiological features as well as distinct responses to stimuli have been observed in astrocytes from different mouse brain regions (Takata and Hirase 2008; Chai et al. 2017; Morel et al. 2017; Itoh et al. 2018; Kostuk et al. 2019; Makarava et al. 2019; Xin et al. 2019; Lozzi et al. 2020). Indeed, astrocyte heterogeneity has been suggested to underlie the regional susceptibility to human diseases (Schober et al. 2022). Therefore, recapitulating astrocyte regional specificity in PSC-derived astrocytes is generally accepted as an important prerequisite.

Several studies have described the generation of regional astrocytes from human embryonic stem cells or induced pluripotent stem cells (iPSCs), including the forebrain (Krencik et al. 2011; Zhou et al. 2016; Tcw et al. 2017; Lin et al. 2018; Bradley et al. 2019, Hedegaard et al. 2020; Peteri et al. 2021), and ventral midbrain (Booth et al. 2019; Barbuti et al. 2020; Crompton et al. 2021; de Rus Jacquet et al. 2021), hindbrain and spinal cord (Roybon et al. 2013; Serio et al. 2013; Holmqvist et al. 2015; Bradley et al. 2019; di Domenico et al. 2019; Yun et al. 2019). The regional identity of these astrocytes is typically evaluated at the stage of early neural progenitors generated via cell type- or region-directed neural patterning protocols, with the assumption that the positional characteristics will be faithfully preserved in the final astrocyte products. However, astrocyte production in vitro involves an extended period of astrocytic fate induction and progenitor expansion using FGF and EGF, and substantial literature has reported alterations in region-specific gene expression and/or neurogenic competence in expanded neural progenitors (Jain et al. 2003; Sun et al. 2008; Koch et al. 2009; Falk et al. 2012). Therefore, better characterization of PSC-derived astrocytes and their lineage-specific features is needed to advance our knowledge of the molecular heterogeneity of human astrocytes.

Using a human iPSC line that allows the tracing of LMX1A expressing ventral midbrain neural progenitors and their differentiated progeny (Cardo et al. 2023), we discovered an unexpected gradual depletion of LMX1A^+^ progenitor progeny during astrocyte induction from a bulk population of ventral midbrain patterned progenitors, despite LMX1A^+^ progenitors being the predominant starting population. However, LMX1A^+^ progenitor-derived astrocytes can be generated if astrocytic induction is initiated from purified LMX1A^+^ progenitors, indicating that the positional constituents of the founding cell population may not be preserved faithfully in the derived astrocytes. Single-cell RNA sequencing (scRNAseq) of astrocytes derived from both parental populations identified distinct transcriptomic signatures, providing a useful resource for the assessment of a defined lineage of human PSC-derived midbrain astrocytes.

## Results

### Depletion of LMX1A^+^ progenitors and/or derivatives in ventral midbrain patterned neural progenitor cultures during astrogenic induction

To investigate whether regionally patterned neural progenitors retain their lineage identity during astrogenic induction and glial progenitor expansion, we made use of the LMX1A-Cre/AAVS1-BFP iPSCs tracer line, in which LMX1A-driven Cre activates BFP expression, thereby enabling tracking of LMX1A^+^ ventral midbrain progenitors and their differentiated progeny (Cardo et al. 2023). We differentiated the LMX1A-Cre/AAVS1-BFP iPSCs towards the ventral midbrain fate following a modified protocol based on Jaeger *et al*. and Kriks et al (Jaeger et al. 2011; Kriks et al. 2011; Figure 1A). Immunocytochemistry of day (d) 19 cultures confirmed a high proportion of cells expressing BFP (96.01±0.42%) and ventral midbrain progenitor markers LMX1A (92.94±0.91%), FOXA2 (94.76±0.57%), and OTX2 (97.82±0.28%; Figure 1B-D with the original images shown in Figure S1A). Most cells (91.26±1.64%) co-expressed LMX1A and FOXA2 (Figure 1B and 1D). At this stage, all BFP^+^ cells stained positive for the pan-neural progenitor marker NESTIN (Figure 1C). We detected a small proportion of cells expressing the midbrain basal plate marker NKX6.1 (2.03±0.47%, Figure 1B-C), whose expression domain in the early developing ventral midbrain partially overlap with that of LMX1A (Andersson et al. 2006). However, few PAX6^+^ cells were present (Figure 1C), which marks the forebrain and lateral midbrain (Duan et al. 2013). Immunocytochemical analysis of FACS-sorted BFP^+^ cells confirmed highly enriched expression of LMX1A (95.51±0.09%) and FOXA2 (95.35±0.14%), and co-expression of both markers (94.68±0.10%; Figure 1B and 1D). In contrast, only a small number of LMX1A^+^ cells (4.96±0.70%) were present in the sorted BFP^-^ population (Figure 1B and1D). These findings provide further support that BFP expression faithfully recapitulates LMX1A expression, and that LMX1A^+^ ventral midbrain progenitors represent the major cell population at d19 (Cardo et al. 2023).

**Figure 1.**
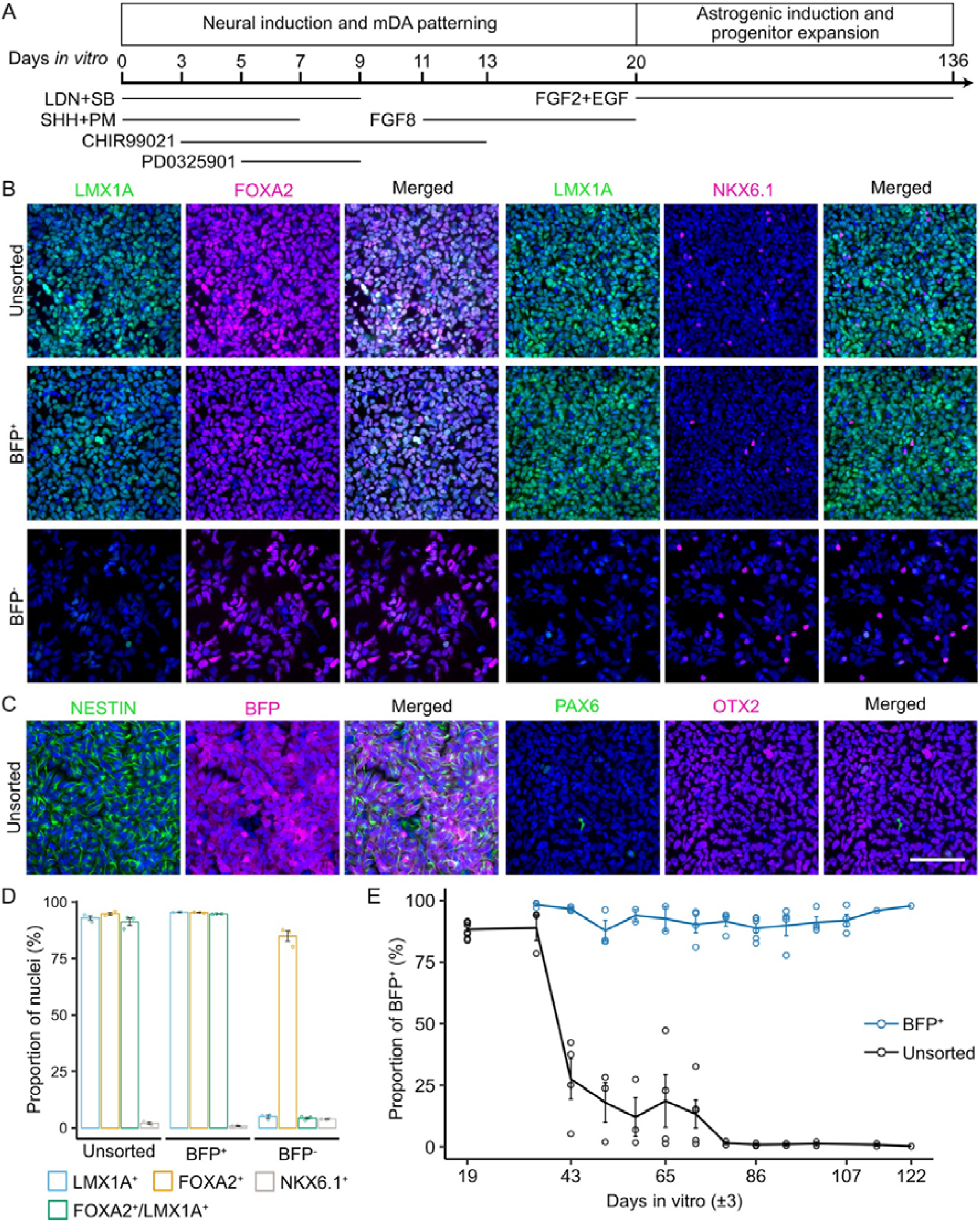
Depletion of LMX1A^+^ progenitors and their derivatives during astrogenic induction in ventral midbrain patterned neural progenitor cultures. A, Schematic diagram of ventral midbrain neural differentiation and astrogenic induction. B-C, Representative view of immunocytochemistry of ventral midbrain neural progenitor markers and other regional markers in d19 unsorted, sorted BFP^+^, and sorted BFP^-^ population. Scale bar represents 100 µm. Images shown were cropped to 300 µm × 300 µm by randomly selecting the region of interest in the nuclei-only channel (uncropped greyscale images are shown in Figure S1A). D, Quantification of marker expression in unsorted, sorted BFP^+^, and sorted BFP^-^ population. Error bars represent the standard error of means (SEM) of three independent experiments. E, Flow cytometry quantification of unsorted and BFP+ population during astrogenic induction and progenitor expansion. Each data point represents one biological replicate. The gating strategy used is shown in Figure S1B.

The d19 cells were then induced to undergo astrogenic switch in media containing FGF2 and EGF with the BFP^+^ cells (LMX1A^+^ ventral midbrain progenitors or their derivatives) monitored by flow cytometry at each weekly passaging (Figure 1A, representative gating strategy is shown in Figure S1B-E). Unexpectedly, we found a dramatic decrease in the BFP^+^ cell proportion from the starting point of 88.20±1.10% to only 27.59±8.28% at d43 and nearly absent as differentiation continued (Figure 1E). We did not observe any evident cell death during culture and replating; thus, the absence of BFP could be either due to the silencing of BFP expression in the derivatives of LMX1A^+^ progenitors or the loss of these cells through growth competition. To address this question, we performed progenitor expansion and astrogenic induction under the same culture conditions as purified d19 BFP^+^ progenitors isolated using fluorescence-activated cell sorting (FACS). Interestingly, the proportion of BFP^+^ cells remained at approximately 90% throughout the astrogenic induction and glial progenitor expansion periods (Figure 1E). This observation demonstrates that BFP expression can be maintained in the derivatives of LMX1A^+^ midbrain progenitors and that the loss of BFP^+^ cells in the unsorted culture is likely due to their growth disadvantage compared to the derivatives of LMX1A^-^ progenitors. Our findings are unexpected and demonstrate that the regional or lineage identity of PSC-derived astrocytic cells should not be assumed based merely on the dominant lineage identity of their cellular origin, given that no in vitro fate specification paradigm is 100% efficient.

### Astrogenic switch occurred earlier in derivatives of LMX1A^+^ midbrain progenitors

Since sorted BFP^+^ (LMX1A^+^ ventral midbrain progenitors or their derivatives) and unsorted astrocytic cultures differ distinctively in BFP expression soon after the initiation of astrogenic induction, for simplicity, these cultures are hereafter referred to as the BFP^+^ and BFP^-^ cultures, respectively. To determine whether the two cell populations behave differently in the process of astrogenic switch, we examined astrocytic marker expression in these cultures at d45 and d98 using immunocytochemistry. SOX9 and NFIA are transcription factors that are crucial for the initiation of astrogenesis and acquisition of astrogenic competence in the developing central nervous system (Stolt et al. 2003; Deneen et al. 2006), whereas CD44 has been identified as an astrocyte-restricted precursor (Liu et al. 2004). We found that all these markers were more abundantly detected in the BFP^+^ cultures (NFIA: 65.89±2.81%; SOX9:57.19±4.25%) than in the FP cultures (NFIA: 4.26±1.28%; SOX9:8.88±1.82%) at d45 (Two-way ANOVA with post-hoc Tukey test, NFIA: p=2.52×10^-5^, SOX9: p=9.21×10^-6^; Figure 2A and 2C with the original images shown in Figure S2). Although the number of NFIA^+^ and SOX9^+^ cells significantly increased in the BFP^-^ cultures by d98 (NFIA: 44.07±4.56% on d98, p=3.58×10^-4^; SOX9:44.28±2.84% on d98, p=1.21×10^-4^), the BFP^+^ cultures still contained more cells expressing NFIA (65.71±4.25%; p=2.06×10^-2^) and SOX9 (73.25±2.12%; p=5.51×10^-4^) than in BFP^-^ cultures (Figure 2A and 2C).

**Figure 2.**
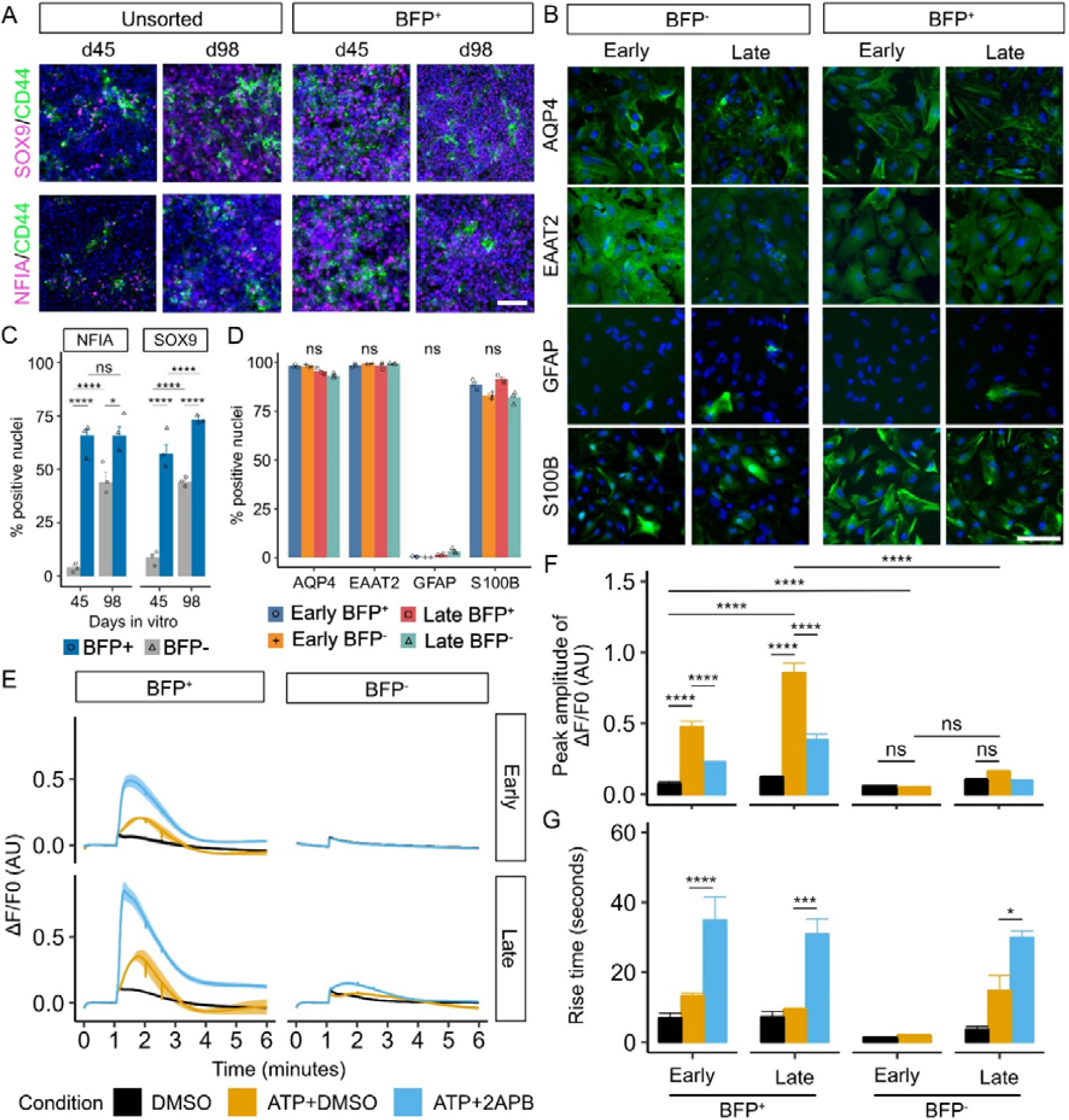
Early astrogenic switch and astrocyte maturation in derivatives of LMX1A^+^ midbrain progenitors. A, Representative view of immunocytochemistry of astrogenic marker expression in BFP^+^ and unsorted progenitors at day 45 and 98. Scale bar represents 100 µm. Images shown were cropped to 462 µm × 462 µm by randomly selecting the region of interest in the nuclei-only channel (uncropped greyscale images are shown in Figure S2). B, Representative view of immunocytochemistry of astrocyte marker expression in early and late, BFP^+^ and BFP^-^ (unsorted) astrocytes. Scale bar represents 100 µm. Images shown were cropped to 300 µm × 300 µm by randomly selecting the region of interest in the nuclei-only channel (uncropped greyscale images are shown in Figure S3). C, Quantification of immunocytochemistry of astrogenic marker in BFP^+^ and BFP^-^ progenitors at day 45 and 98 shown in Panel A. Error bars represent the standard error of means (SEM) of three independent experiments. Two-way ANOVA was performed to compare between lineages (NFIA: p=5.389×10^-6^, df=1, effect size=3.62; SOX9: p=1.96×10^-6^, df=1, effect size=4.77) and days of differentiation (NFIA: p=7.82×10^-5^, df=1, effect size=1.99; SOX9: p=2.62×10^5^, df=1, effect size=2.99) D, Quantification of immunocytochemistry of astrocyte marker expression in astrocytes. Error bars represent SEM of three independent experiments. Kruskal-Wallis test results following Bonferroni correction are shown on the top of the figure (AQP4: p.adjust=0.12, df=3, H=8.95; EAAT2: p.adjust=1.00, df=3, H=0.95; GFAP: p.adjust=0.06, df=3, H=10.38; S100B: p.adjust=0.11, df=3, H=9.05). E, Averaged trace of ATP-induced Ca^2+^ response assayed using FLIPR. Drugs or DMSO were applied at 1 minute of the assay. The line represents the average fluorescence change (ΔF/F0) in at least three independent experiments each with at least three replicate^Δ^wells. The shaded area represents the SEM across at least three independent experiments. F, Quantitative comparison of the peak amplitude of ATP-induced Ca^2+^ response among conditions (two-way ANOVA, p<2.2×10^-16^, df=2, effect size=2.54) and samples (p=2.87×10^-14^, df=3, effect size=2.17). Error bars represent the SEM across at least independent experiments. G: Quantitative comparison of the rise time of ATP-induced Ca^2+^ response among conditions (two-way ANOVA, p=2.19×10^-13^, df=2, effect size =1.958) and samples (p=0.064, df=3, effect size=0.76). Intergroup comparison was performed using post-hoc Tukey test. Error bars represent the SEM across at least three independent experiments. (****: p<0.0001, ***: p<0.001, **: p<0.01, *: p<0.05, ns: not significant).

To investigate whether the temporal difference in the astrocytic switch between the BFP^+^ and BFP^-^ cultures affects the maturation and functionality of the derived astrocytes, we initiated astrocyte terminal differentiation by exposing the BFP^+^ and BFP^-^ astrocyte precursors to CNTF and BMP4 (Krencik et al. 2011; Bradley et al. 2019) from d87 (referred to as early astrocytes) and d136 (late astrocytes). Both BFP^+^ and BFP^-^ cultures exhibited a similar expression profile of classic astrocyte markers, including AQP4, EAAT2, and S100B, but few GFAP^+^ cells at both time points (Figure 2B and 2D with the original images shown in Figure S3). As a reference, we also generated neural progenitors without employing any patterning cues and induced astrogenic switch and astrocyte differentiation from these non-patterned neural progenitors (Figure S4A-B). We found that, while the astrocyte cultures derived from the non-patterned progenitors contained a similar proportion of cells expressing AQP4, EAAT2 and S100B compared to the BFP^+^ and BFP^-^ astrocyte cultures, there are more GFAP^+^ cells in the non-patterned astrocyte preparations (17.89±5.4%, Figure S4A-B).

Functional astrocytes exhibit transient calcium (Ca^2+^) spikes upon chemical stimulation such as ATP (Zhang et al. 2016). Using a FLIPR Ca^2+^ release assay, we observed a sharp increase in the intracellular Ca^2+^ concentration upon ATP administration in both the early and late BFP^+^ astrocyte populations (early BFP^+^: p=5.37 10^-12^; late BFP^+^: p<2.2×10^-16^; Figure 2E-F). ATP-induced Ca2^+^ release is partially mediated by inositol trisphosphate. Indeed, addition of an inositol trisphosphate receptor antagonist 2-aminoethoxydiphenylborate (2-APB) reduced the amplitude (early BFP^+^: p=5.88×10^-5^; late BFP^+^: p=9.22×10^-10^; Figure 2F) and rise time (early BFP^+^: p=2.63×10^-5^; late BFP^+^: p=3.59×10^-4^; late BFP^-^: p=0.028; Figure 2G) of ATP-induced Ca2^+^ response in both the early and late BFP^+^ astrocytes. Interestingly, early BFP^+^ astrocytes had a significantly lower peak amplitude than that observed in late BFP^+^ astrocytes (p=6.92×10^-9^; Figure 2F), despite their similar levels of astrocyte marker expression, suggesting a difference in maturity at the functional level. Early and late BFP^-^ astrocytes exhibited a similar profile of time-dependent increase in the amplitude of ATP-induced Ca^2+^ response, but did not reach statistical significance (Figure 2E-F). However, late BFP^-^ astrocytes showed a significantly lower peak amplitude than late BFP^+^ astrocytes (p<2.2×10^-16^; Figure 2E-F).

Taken together, our data demonstrated that astrogenesis occurred earlier in BFP+ cultures than in BFP-cells. This temporal difference is also reflected in the functional maturity of the derived astrocytes despite a similar expression profile of classic astroglial markers.

### Single cell RNA sequencing confirms the authenticity of PSC-derived astrocytes

To further characterize the PSC-derived astrocytes, we performed full-length scRNAseq on early and late BFP^+^ and BFP^-^ astrocytes using the iCELL8 platform and SMART-seq technology, with non-patterned astrocytes derived from the LMX1A-Cre/AAVS1-BFP tracer line as a control. A sample of iPSC-derived neurons was included to facilitate the downstream cell type identification. We profiled FACS purified astrocytes expressing CD49f as well as unsorted cultures for comparison of all three astrocyte populations (Barbar et al. 2020); Figure S4C-H). After stringent filtering (Figure S5A-C, see Methods and Materials on filtering), we obtained 17478 protein-coding genes in 1786 qualifying cells, with an average of 6326 protein-coding genes detected per cell.

Unsupervised Louvain clustering identified 12 cell clusters (Figure 3A). Cells were clustered mainly based on sample type (astrocytes and neurons; Figure S6A) and the estimated cell cycle phase (Figure S6B), whereas sorted CD49f^+^ and unsorted astrocytes were largely intermingled together (Figure S6A). Clusters 0, 1, 4, 5 and 11 consisted of predominantly BFP^+^ astrocytes, each with varying contribution from the ‘early’ or ‘late’ astrocyte samples. Clusters 2, 6, and 7 were dominated by BFP-astrocytes, with the majority of the ‘early’ astrocyte samples in cluster 2. Cells in clusters 8,9 and 10 were primarily non-patterned astrocytes, whereas cluster 3 came from the neuronal sample (Figure 3B-C). Consistent with immunocytochemistry, *TagBFP* transcripts were detected at a higher level in BFP^+^ astrocyte samples than in BFP^-^ and NP samples, while *TagBFP* expression was negligible in neuronal samples derived from an iPSC line without the BFP transgene (Figure S6C-D). Using a set of known astrocyte and neuronal signature genes (Figure 3D), we identified cells in clusters 0, 1, and 5-11 as astrocytes (Figure 3D), which were enriched in SOX9, NFIA, NFIB, HES1, S100A13, S100A16, EGFR, CD44, and GJA1 expression (Figure 3D). These transcripts were also detected at high levels in clusters 2 and 4, which were mostly estimated to be in the cell cycle phases G2, M, and S (Figure S6B). In addition, clusters 2 and 4 showed high levels of proliferation-related transcripts, such as *TOP2A, MKI67, CDK1*, and *AURKA* (Figure 3D) and were thus defined as astrocyte precursors. In contrast, Cluster 3 contained mostly cells from the neuronal sample (Figure 3A and Figure S6A) and expressed high levels of genes closely related to neuronal structure and function (such as *STMN2*, *SYT1*, *DCX*, *MAPT*, and *SNAP25*; Figure 3D). We did not detect transcripts indicative of endoderm (*GATA4*), mesoderm (*TBXT* and *TBX6*), or oligodendrocyte progenitors (*SOX10* and *PDGFRA*) in any of these clusters (Figure S6C).

**Figure 3.**
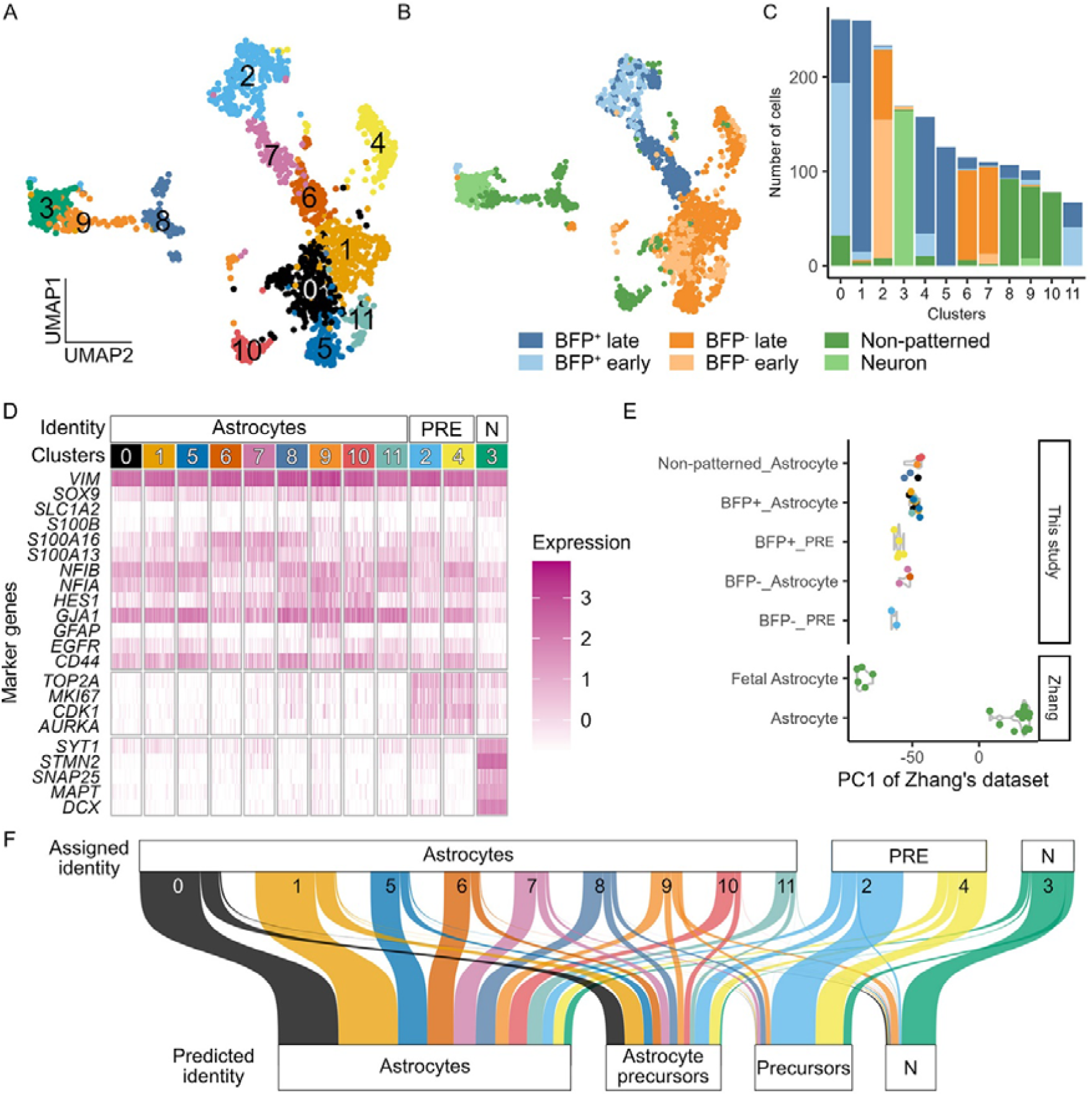
Single cell RNA sequencing confirms the authenticity of PSC-derived astrocytes. A-B , Uniform manifold approximation and projection plot of unbiased clustering, coloured by clusters or sample group. C, Number of cells from different sample groups in each clusters. D, Heatmap of the normalised expression of selected markers in different clusters. The assigned identity to each cluster is shown at the top of the plot. E, Principal component projection analysis of pseudobulk astrocyte data onto a reference principal component axis of astrocyte maturity (Zhang et al. 2016; Hedegaard et al. 2020). Each dot represents a pseudobulk sample of one independent sample from each clusters. F, Sankey plot summarising the result of reference mapping of cells in different clusters to eight published reference human brain scRNAseq datasets. The thickness of the thread is proportional to the number of cells mapped to the same identity in the reference datasets (predicted identity). Detailed results of referencing mapping to each reference datasets are shown in Figure S7A-H and prediction score shown in Figure S7I. (PRE: precursors; N: neurons).

Next, we examined the developmental status of astrocytes by pseudobulk analysis using published bulk RNA-seq datasets from human fetal and postmortem astrocytes (Zhang et al. 2016). As expected, the astrocyte precursors (clusters 2 and 4) were projected to be less mature than the fetal astrocytes (Figure 3E). The BFP^+^ astrocyte dominant clusters (0, 1, 5, and 11) and non-patterned astrocyte clusters (8, 9, and 10) were shown to be more developmentally advanced than the BFP^-^ astrocyte clusters (6 and 7). Thus, the pseudobulk analysis provides independent support to the functional assays suggesting the relative maturity of BFP^+^ and BFP^-^ astrocytes (Figure 2).

To determine the authenticity of these PSC-derived astrocytes, we mapped our data to five published scRNAseq datasets obtained from human fetal and adult brains using Seurat integration (La Manno et al. 2016; Sloan et al. 2017; Zhong et al. 2018; Polioudakis et al. 2019; Agarwal et al. 2020; Fan et al. 2020; Bhaduri et al. 2021; Eze et al. 2021). We found that cells in clusters annotated as astrocytes (clusters 0, 1, and 5-11) were predominantly mapped to the reference astrocyte or astrocyte precursor populations with high confidence (prediction score > 0.5; Figure 3F and Figure S7). In contrast, neuronal cluster 3 was mapped to neurons in the fetal reference datasets, while the astrocyte precursor clusters (2 and 4) were mapped to progenitor populations in the fetal reference datasets (Figure 3F and Figure S7). These findings demonstrate that iPSC-derived astrocytes closely resemble those in the human brain.

### Distinct transcriptome fingerprints of LMX1A^+^ midbrain progenitor-derived astrocytes

Significant advances have been made recently in the understanding of the molecular profiles of midbrain dopamine neurons. However, our knowledge of midbrain astrocytes in this regard remains limited and does not inform the anatomic or lineage origin of the cells. In this regard, BFP^+^ astrocytes provide a unique resource for determing the transcriptomic characteristics of human astrocytes derived from the LMX1A^+^ lineage of ventral midbrain patterned progenitors. By performing pairwise differential gene expression (details described in Methods and Materials), we identified 1153 genes differentially expressed (DEGs; adjusted p values less than 0.05 and log2 fold change over 0.25) in BFP^+^ astrocytes when compared to either BFP^-^ or non-patterned astrocyte populations (Supplementary Data 1). Of these, 287 were unique to BFP^+^ astrocytes (BFP^+^ enriched, Figure 4A), including genes associated with midbrain dopamine neuron development such as *SULF1*, *LMO3*, *NELL2*, and *RCAN2* (Figure 4B) (Strelau et al. 2000; La Manno et al. 2016; Bifsha et al. 2017; Ahmed et al. 2021). 55 of the 287 BFP^+^ DEGs, including *SULF1*, *NELL2*, *RCAN2*, and *RGS5,* were confirmed to be significantly preferentially expressed in midbrain astrocytes in vivo (adjusted p-values less than 0.05) by analysing the integration of five human brain datasets (Figure S8A-B; La Manno et al. 2016; Zhong et al. 2019; Fan et al. 2020; Eze et al. 2021). Another 87 BFP^+^ DEGs, such as *SLC7A3*, *SIX6*, and *LMO3*, were also detected at higher levels in midbrain astrocytes than in forebrain astrocytes, although their expression levels were not significant (Figure S8A-B). Moreover, *LMX1A* and *FOXA2*, which have been used to evaluate PSC-derived midbrain astrocytes in previous studies, were not detected in BFP^+^ astrocytes (Figure S6D, S8C).

**Figure 4.**
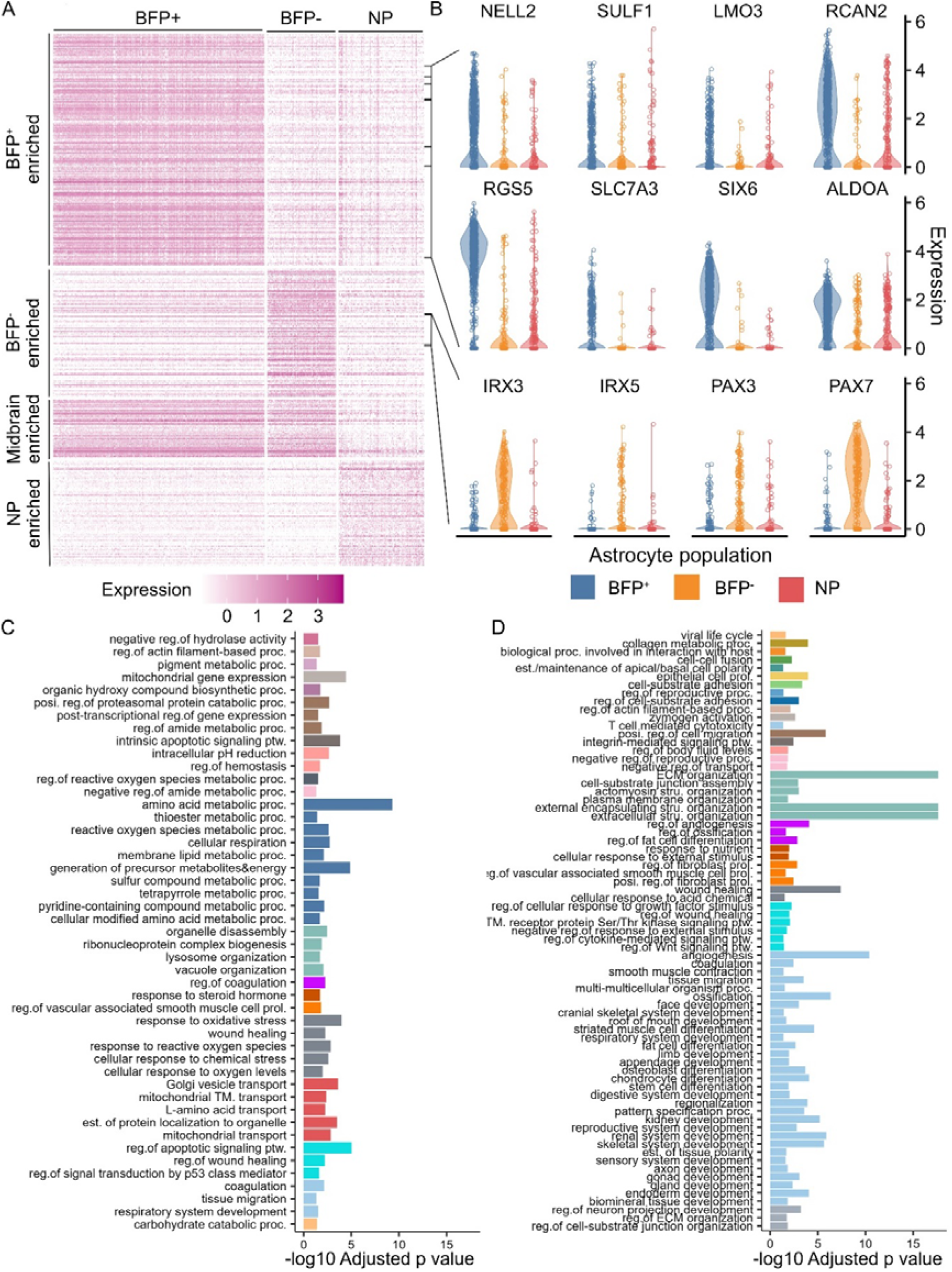
Distinct transcriptome fingerprints of astrocytes derived from LMX1A^+^ ventral midbrain progenitors. A, Heatmap of the normalised expression of population-specific genes in different populations of astrocytes. B, Violin plots of the normalised expression of selected candidate markers for BFP^+^, BFP^-^, and non-patterned (NP) astrocytes. C-D, Representative GO terms significantly enriched in BFP^+^ (C) and BFP^-^ (D) enriched genes. Semantically similar representative terms were shown with the same colour.

We also identified 159 DEGs enriched only in BFP^-^ astrocytes (BFP^-^ enriched, Figure 4A and Supplementary Data 1), including those known to be expressed in the ventrolateral-dorsal domain of the midbrain and hindbrain, such as *IRX3*, *IRX5*, *PAX3*, and *PAX7* (Figure 4B) (Houweling et al. 2001; Matsunaga et al. 2001). Differential expression of PAX3 and PAX7 in BFP astrocytes was confirmed at the protein level by immunocytochemistry of d136 BFP^+^ and BFP^-^ astrocyte cultures, whereas their expression in the human midbrain was validated by published datasets in silico (Figure S8). This transcription profile supports the notion that BFP astrocytes are descendants of initial minor populations of lateral midbrain progenitors. Moreover, 72 DEGs were shared by BFP^+^ and BFP^-^ astrocytes compared to non-patterned astrocytes (midbrain enriched, Figure 4A and Supplementary Data 1). This set of genes included *NR2F1*, *NR2F2*, *ZEB2*, *KCNJ6*, and *SRPX* (Figure 4B), which have been reported to be signatures of mouse midbrain astrocytes (Endo et al. 2022). Together, our findings provide a new entry into the transcriptomic characteristics of midbrain astrocytes, specifically a gene expression map of the LMX1A^+^ midbrain progenitor-derived human astrocyte lineage.

Gene ontology (GO) enrichment analysis was performed on the 1153 DEGs enriched in BFP^+^ astrocytes (Supplementary Data 2). The significantly enriched GO terms were mainly related to various aspects of metabolism, stress response, biosynthesis, lysosomal activity, and cellular respiration (Figure 4C and Supplementary Data 3). These biological processes have previously been shown to be disrupted by several mutations that cause familial Parkinson’s disease (di Domenico et al. 2019; Barbuti et al. 2020; Sonninen et al. 2020). In contrast, the GO terms associated with 530 BFP^-^ astrocyte enriched DEGs were mostly related to the formation of the extracellular matrix and tissue development (Figure 4D and Supplementary Data 2-3). The differential enrichment of GO terms implies functional differences between BFP^+^ and BFP^-^ astrocytes and supports the need for generating regional specific astrocytes for disease modelling.

## Discussion

Despite the general belief that recapitulating astrocyte lineage heterogeneity is necessary for stem cell-based disease modelling and cell transplantation, the extent of astrocyte heterogeneity in different brain regions, their anatomical origins, and associated molecular signatures remain largely elusive. This knowledge gap limits the end-point characterization of stem cell-derived astrocytes; hence, reliance on the dominant regional characteristics of the initial neural progenitor populations, with the assumption that the lineage representation of the progenitors is preserved in the derived astrocytes. By harnessing an LMX1A based lineage tracing human iPSC line, we discovered unexpected negative selection against derivatives of LMX1A^+^ midbrain-patterned progenitors during astrocyte induction and progenitor expansion, highlighting the need for careful characterization of PSC-derived astrocytes and reinforce the need for a deeper understanding of the molecular landscape of astrocytes in different regions of the human brain.

Most neural progenitors used for astrocyte differentiation in this study coexpressed LMX1A and FOXA2. In the ventral midbrain, the combinatorial expression of these transcription factors defines the dopaminergic neural progenitors (Failli et al. 2002; Andersson et al. 2006). We found that astrocytes derived from LMX1A^+^ progenitors could only be obtained if LMX1A^+^ cells were purified prior to astrocyte differentiation. In contrast, astrocytes derived from bulk midbrain-patterned progenitors exhibit transcriptomic profiles of the lateral-dorsal midbrain despite LMX1A^+^ progenitors being the predominant starting population. Our findings demonstrate that the lineage composition of parent progenitors was not preserved during astrocyte induction or progenitor expansion. FGF is the most commonly used inductive molecule for astrocyte differentiation from stem cells (Chandrasekaran et al. 2016). However, it is evident that FGF-expanded neural progenitors, originating either from the brain or neutralized PSCs, exhibit restricted regional competence and positional gene expression. For example, bulk-expanded human ventral midbrain neural progenitors (Jain et al. 2003), fetal forebrain or spinal cord derived neural stem (NS) cells only give rise to GABAergic neurons (Sun et al. 2008), and lt-NES cells display an anterior hindbrain-like positional profile (Falk et al. 2012), while their antecedents, PSC-derived neural rosettes and early passage derivatives, express anterior forebrain markers (Koch et al. 2009). It is unclear whether this is due to the deregulation of the original patterning at the level of gene expression or the loss of the associated cell population (Gabay et al. 2003). In this study, because BFP^+^ astrocytes can be generated under the same culture conditions as purified LMX1A^+^ progenitors, we reasoned that the loss of their derivatives in unsorted cultures was possibly due to their differential growth capacity.

Our study highlights the need for a careful assessment of the positional identity of in vitro-derived astrocytes. A common practice in this regard is to confirm the regional identity of founder progenitors following fate-directed neural induction, with the assumption that the dominant positional features are maintained by astrocyte progeny (Krencik et al. 2011). This strategy is, at least partly, dictated by our limited knowledge of the gene expression signatures of regional and/or lineage-specific astrocytes. Hence, endpoint evaluation of PSC-derived astrocytes often relies on region-specific markers defined in the brain during the neurogenic period. For example, LMX1A and FOXA2 expression has been used as criteria for midbrain astrocytes in previous studies (Barbuti et al. 2020; Crompton et al. 2023). However, scRNA-seq of the human fetal ventral midbrain and adult substantia nigra has revealed negligible expression of these transcripts in astrocytes (La Manno et al. 2016; Agarwal et al. 2020; Kamath et al. 2022). Consistent with these findings, we did not detect LMX1A or FOXA2 in BFP^+^ or BFP ^−^ astrocytes. However, our analysis identified new positive and negative markers that can be used to validate astrocytes derived from the LMX1A^+^ lineage of ventral midbrain progenitors. Future work will benefit from in vivo validation of these putative lineage-specific markers, as a mouse analogous tracer line is available, whereas lineage tracking is not possible in humans.

In addition to the distinct transcriptomic profiles, BFP^+^ and BFP astrocytes may also be functionally different. Astrocytes generated from progenitors broadly patterned to the dorsal forebrain, ventral forebrain, and spinal cord have been shown to exhibit different GO enrichment profiles as well as different physiological and functional properties (Bradley et al. 2019). In comparison to BFP^-^ and non-patterned astrocytes, the current study revealed that GO terms enriched in BFP^+^ astrocytes, which originated from the same progenitor giving rise to midbrain dopaminergic neurons, were closely related to various biological processes disrupted in astrocytes carrying familial Parkinson’s disease mutations (di Domenico et al. 2019; Barbuti et al. 2020; Sonninen et al. 2020). Such a distinct enrichment profile suggests that BFP^+^ astrocytes may be functionally adapted to support midbrain dopaminergic neurons compared with BFP^−^ and non-patterned astrocytes. Indeed, astrocytes isolated from the ventral midbrain have been reported to exhibit stronger neurotrophic effects and the ability to reduce α-synuclein aggregation in the midbrain than cortical astrocytes in cellular and mouse models of PD (Kostuk et al. 2019; Yang et al. 2022), highlighting the importance of understanding astrocyte heterogeneity in iPSC disease modelling.

In conclusion, this study provides further evidence for the regional diversity of astrocytes and identifies a set of midbrain-enriched genes. Crucially, the transcriptomic fingerprint of human astrocytes derived from LMX1A-expressing midbrain progenitors reported here offers a much-needed resource for assessing the authenticity of stem cell-derived astrocytes in studies of Parkinson’s disease.

## Methods and Materials

### Stem cell culture and astrocyte differentiation

KOLF2 human iPSCs were maintained in E8 flex media (ThermoFisher) and manually dissociated using Gentle Cell Dissociation Reagent (STEMCELL Technologies) as previously described (Cardo et al. 2023). Astrocytes were differentiated using a three-stage stepwise strategy consisting neural induction and regional patterning, astrogenic switch and progenitor expansion, and astrocyte terminal differentiation. LMX1A^+^ ventral midbrain progenitors were generated as previously described (Cardo et al. 2023). At day 19, cells were replated as single cells onto poly-D-lysine-laminin-coated plates at 1×10^6^ cells/cm^2^ for astrogenic switch and progenitor expansion in N2B27 media supplemented with 10 ng/mL FGF2 (Peprotech) and 10 ng/mL Human EGF (Peprotech) and replated every 6-8 days. For astrocyte terminal differentiation, expanded neural progenitors were re-plated at a density of 3×10^4^ cells/cm^2^ in expansion media and 24 hours later switched to N2B27 supplemented with 10 ng/mL human recombinant CNTF (Peprotech) and 10 ng/mL human recombinant BMP4 (Peprotech) for 7 days followed by media containing CNTF alone for another 13 days. 10 µM Y-27632 was used for 24 hours before and after each replating. The protocol for generating non-patterned astrocytes was the same as for LMX1A^+^ ventral midbrain-derived astrocyte except the neural progenitors were derived with duo-SMAD inhibitors only without ventral patterning reagents.

### Flow cytometry analysis and cell isolation

Cells were dissociated in Accutase as described above and washed twice with DPBS by centrifugation for 5 minutes at 200 rcf. For evaluating BFP expression, dissociated cells were resuspended in 0.5 mM EDTA in DPBS (Sigma-Aldrich) and analysed on a BD LSRFortessa cell analyser (BD Biosciences). For purifying BFP^+^ cells, dissociated cells were resuspended in the same cell culture media.

Background autofluorescence was compensated for using KOLF2 parental cell line at a similar stage of differentiation to define BFP^-^ gating. For purifying CD49f^+^ astrocytes, dissociated cells were stained with Alexa Fluor 647-conjugated rat anti-CD49f antibody (5% v/v in a 100 µL reaction; BD Biosciences) for 25 minutes at 37°C on an orbital shaker at 200 rcf and resuspended in DPBS containing 0.5% bovine serum albumin and 50 units/mL DNase I (Sigma Aldrich). Background autofluorescence was compensated for using KOLF2 parental cell line at a similar stage of differentiation to define BFP^-^ gating and unstained astrocytes to define CD49f^-^ gating. Cell sorting was performed on a BD FACSAria III (BD Biosciences) using an 80 µm nozzle. Sorted cells were collected in the same media as for resuspension. Flow cytometry data were analysed in FlowJo v10.8.1 (BD Biosciences) as shown in Figure S1B-E. Briefly, non-debris events were selected using the eclipse gates on dot graphs of SSC-A versus FSC-A. Singlet events were sequentially gated using polygonal gates on dot graphs of FSC-H versus FSC-A and SSC-H versus SSC-A by selecting the events in the diagonal region. The positive and negative gates in the fluorescence channel were set as bifurcate gates at a minimum of 99.9% percentile (usually at 99.99% percentile) on the histogram of the fluorescence intensity of the negative control sample of the same flow cytometry experiment and applied to all samples of the same flow cytometry experiment.

### FLIPR calcium assay

Astrocytes were plated on day 10 of astrocyte terminal differentiation and cultured according to the standard protocol until day 20. On the day of recording, equal volume of FLIPR Calcium 6 Assay buffer was added to cells without replacing or washing and cells were incubated for 2 hours at 37°C. Drug assay buffers were prepared in HBSS with Ca^2+^ and Mg^2+^ (Gibco) with 20 mM HEPES buffer (Gibco) at 5X concentration. After incubation, cells were imaged on FLIPR Penta system (Molecular Devices) at a frequency of 2 Hz for 1 minute prior to injection. 25 µL of the drug assay buffers were dispensed at a speed of 25µL/second and images captured at a frequency of 2 Hz for a further 5 minutes. Raw fluorescence intensity data were exported and analysed in R 4.3.0 using a custom script to identify the peak response. The average baseline fluorescence intensity for each well was calculated by averaging the raw fluorescence intensity measured 60 seconds prior to drug application. The normalised change in fluorescence intensity above the baseline (ΔF/F0) for each well, at a given timepoint, was calculated using the following formula:

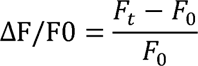

where F_t_ is the fluorescence intensity at timepoint t, F_0_ is the average fluorescence intensity of the baseline period (60 seconds prior to drug application).

### Immunocytochemistry

Cultures were fixed with 3.7% PFA for 15-20 min at 4 °C. For nuclear antigen detection, an additional fixation with methanol gradient was performed, which include 5 mins each in 33% and 66% methanol at room temperature followed by 100% methanol for 20 min at -20°C. Cultures were then returned to PBST via inverse 21 gradient and were then permeabilized with three 10-minute washes in 0.3% Triton-X-100 in PBS (PBS-T) and then blocked in PBS-T containing 1% BSA and 3% donkey serum. Cells were incubated with primary antibodies in blocking solution overnight at 4°C. Following three PBS-T washes, Alexa-Fluor secondary antibodies (Thermo Fisher Scientific) were added at 1:1000 PBS-T for 1 hour at ambient temperature in the dark. Three PBS-T washes were then performed that included once with DAPI (Molecular Probes). Images were taken on a Leica DMI6000B inverted microscope. Quantification was carried out in Cell Profiler (Stirling et al. 2021) or manually using ImageJ (Schindelin et al. 2012) by examining at least four randomly selected fields from three independent experiments. The antibodies used are provided in the Supplementary Table 1. Representative images shown in main figures were cropped by randomly selecting the region of interest in the DAPI-stained channel only, with the original unedited images shown in Figure S1A, S2 and S3.

### Single-cell RNA-sequencing

Cells were dissociated with Accutase with 10 units/mL of papain (Sigma-Aldrich) for 10 minutes at 37°C and resuspended in 0.5% bovine serum albumin (Sigma Aldrich) with 50 units/mL DNase I in DPBS without calcium or magnesium (Gibco) and passed through a cell strainer with 35 µm mesh. Cells were stained with 1 µM SYTO16 (Invitrogen) and 0.08% (v/v) propidium iodide (part of the Invitrogen ReadyProbes™ Cell Viability Imaging Kit, Blue/Red) for 20 minutes on ice and then dispensed into the nano-well plates of the ICELL8® cx Single-Cell System (Takara). Wells containing single viable cells were automatically selected using the ICELL8 cx CellSelect v2.5 Software (Takara) with the green and not red logic. Manual triage was performed to recover additional candidate wells that contain viable single cells. The library for sequencing was then prepared using the SMART-Seq ICELL8 application kit (Takara) following the manufacturer’s recommended protocol. Next-generation sequencing was performed using the NovaSeq6000 and the Xp Workflow on a S4 flow cell for 200 cycles of pair-end sequencing.

### Single-cell RNA-sequencing analysis

Using the Cogent NGS Analysis Pipeline software v 1.5.1 (Takara), FASTQ files containing all indices for each chip were demultiplexed to FASTQ files containing one index per file. Adaptor sequences were removed using cutadapt 3.2 with the 22 following settings: -m 15 --trim-n --max-n 0.7 -q 20. Trimmed FASTQ files were aligned to the *Homo sapiens* GRCh38.106 primary assembly with the BFP reporter gene attached to the end of the genome, using STAR 2.7.9a (Dobin et al. 2013) with the following settings: --outSAMtype BAM Unsorted --quantMode TranscriptomeSAM --outReadsUnmapped Fastx --outSAMstrandField intronMotif --chimSegmentMin 12 --chimJunctionOverhangMin 8 --chimOutJunctionFormat 1 –alignSJDBoverhangMin 10 --alignMatesGapMax 100000 --alignIntronMax 100000 -- alignSJstitchMismatchNmax 5 -1 5 5 --chimMultimapScoreRange 3 -- chimScoreJunctionNonGTAG -4 --chimMultimapNmax 20 -- chimNonchimScoreDropMin 10 --peOverlapNbasesMin 12 --peOverlapMMp 0.1 – alignInsertionFlush Right --alignSplicedMateMapLminOverLmate 0 -- alignSplicedMateMapLmin 30. After alignment, gene-level quantification was performed using featureCounts from subread 2.0.0 (Liao et al. 2013) with the following settings: -t exon --primary -R CORE -F GTF -Q 0 -B -g gene_id. The count matrix of each index was combined in R 4.2.0 (Team 2023).

All downstream analysis was performed in R 4.3.0 using Seurat 4.3.0 (Stuart et al. 2019). Gene-level filtering was applied by including only protein-coding genes with at least five total counts across all cells and being expressed in at least 1% of all cells. Poor quality cells were then identified using the *is.outlier* function from the scater 1.28.0 (McCarthy et al. 2017). Poor quality cells were defined as having a high percentage of mitochondrial gene count, or high or low the total number of genes detected, or high total gene counts. The thresholds of each metrics for each sample were determined as twice the median absolute deviation of the sample. Raw gene counts were log normalised with a scale.factor setting of 1×10^5^. Data from the two batches of experiments were integrated using the *FindIntegrationAnchors* and *IntegrateData* based on the common top 2000 highly variable genes and the first 30 dimensions of principal components. The percentage of mitochondrial gene count and total gene count were regressed out using the *ScaleData* function. Principal component analysis (PCA) was performed on the top 2000 high variable genes and the number of principal components used for uniform manifold approximation and projection (UMAP) was determined using the JackStraw method (Chung and Storey 2015). UMAP and unbiased Louvain clustering was performed on the first 33 principal components. Pairwise differential gene expression analysis was performed using the MAST method (Finak et al. 2015) with “Chip” as the latent variable. Gene ontology enrichment analysis was performed using *enrichGO* function in the clusterProfiler 4.10.0 package (Stirling et al. 2021) with all genes in the filtered dataset as the background. GO term database was downloaded using the org.Hs.eg.db 3.18.0 package (Carlson 2019). Revigo v1.8.1 was used to group the representative GO terms based on semantic similarity using a size setting of 0.5, Homo sapiens database, and SimRel method for semantic similarity (Schlicker et al. 2006; Supek et al. 2011).

Published datasets were downloaded from NCBI’s Gene Expression Omnibus (Clough and Barrett 2016) and processed in R 4.2.0. Gene level filtering was performed by retaining only protein-coding genes with more than five total counts across all cells. Gene counts were normalised using the NormalizeData function (scale.factor settings are listed in Supplementary Table 2). PCA was performed based on the top 2000 highly variable genes to obtain the first 50 PCs. Visual inspection of the elbow plot was used to determine the number of PCs for downstream analysis. Batch effect between subjects was evaluated on the two-dimensional PC2∼PC1 plot. Where inter-subject batch effect was observed, Harmony integration was performed based on the PCs selected in the previous step (Supplementary Table 2). UMAP was performed based on either the PCA or Harmony reduction (using the top 30 dimensions), and Louvain clustering was performed (settings shown in Supplementary Table 2). Cluster identities were verified against the reported annotation where possible. For datasets without detailed annotation published or astrocyte lineage reported (Supplementary Table 2), reannotation was performed based on the expression of known markers (Figure S7A-E). Reference mapping was performed using *FindTransferAnchors* and *TransferData* function in Seurat based on the first 30 dimensions of either the PCA or Harmony loadings of the reference dataset.

### Statistical analyses

All data were collected from at least three independent experiments and presented as mean ± standard error of means unless otherwise specified. Data were tested for normality with the Shapiro-Wilk test and for equal variance with Levene test before performing statistical analyses by two-way ANOVA with post-hoc Tukey test for multiple comparisons where relevant. Kruskal-Wallis test with post-hoc Dunn’s test for pairwise comparison was used where parametric test was not suitable. Effect size was calculated as Cohen’s f for ANOVA or eta squared based on the H-statistic for Kruskal-Wallis test. All statistical tests were performed in R4.3.0.

## Acknowledgements

We would like to thank Mark Bishop and Joanne Morgan for conducting FACS and next generation sequencing, respectively. We thank Kathryn Peall and Laura Abram for providing iPSC-derived neurons for scRNAseq. We also thank the support of the Supercomputing Wales project, which is part-funded by the European Regional Development Fund (ERDF) via the Welsh Government.

## Author Contribution

Z.L. and M.L. conceived the study and designed the experiments. Z.L. performed all cell experiments and analysed all data. L.C. and M.L. designed the lineage tracing system, and L.C. generated the cell line with assistance from Z.L. M.R. performed iCELL8 library preparation. J.M.S., F.W., and V.V. provided guidance and contributed to general discussions on scRNAseq data analysis. C.W. contributed to the design of scRNAseq and general discussions throughout the work. Z.L. and M.L wrote the paper. All authors edited and approved the paper.

## Data Availability

The sequencing data discussed in this publication have been deposited in NCBI’s Gene Expression Omnibus (Clough and Barrett 2016) and are accessible through a GEO Series accession number GSE252624.

## Supplementary Information

**Figure S1.**
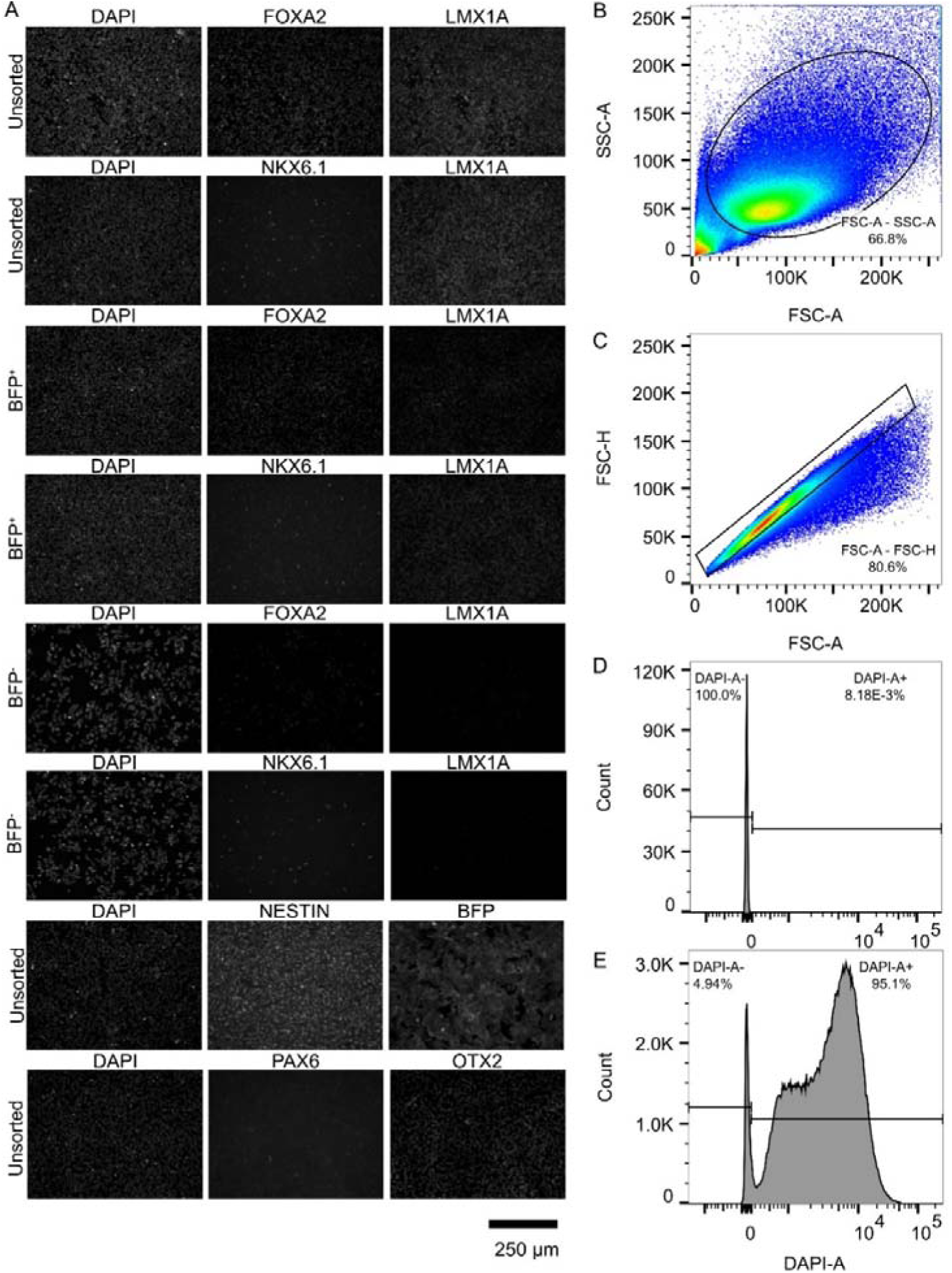
Original images of immunocytochemistry of d19 progenitors and gating strategy of BFP flow cytometry analysis. A, the original images shown in Figure 1B-C. The gating strategy used for BFP flow cytometry analysis are shown in B-E and described in Methods and Materials. B, scatter plot of SSC-A versus FSC-A. C, scatter plot of FSC-H versus FSC-A. D-E, histogram of BFP fluorescence in the negative control (D) and BFP-expressing (E) samples.

**Figure S2.**
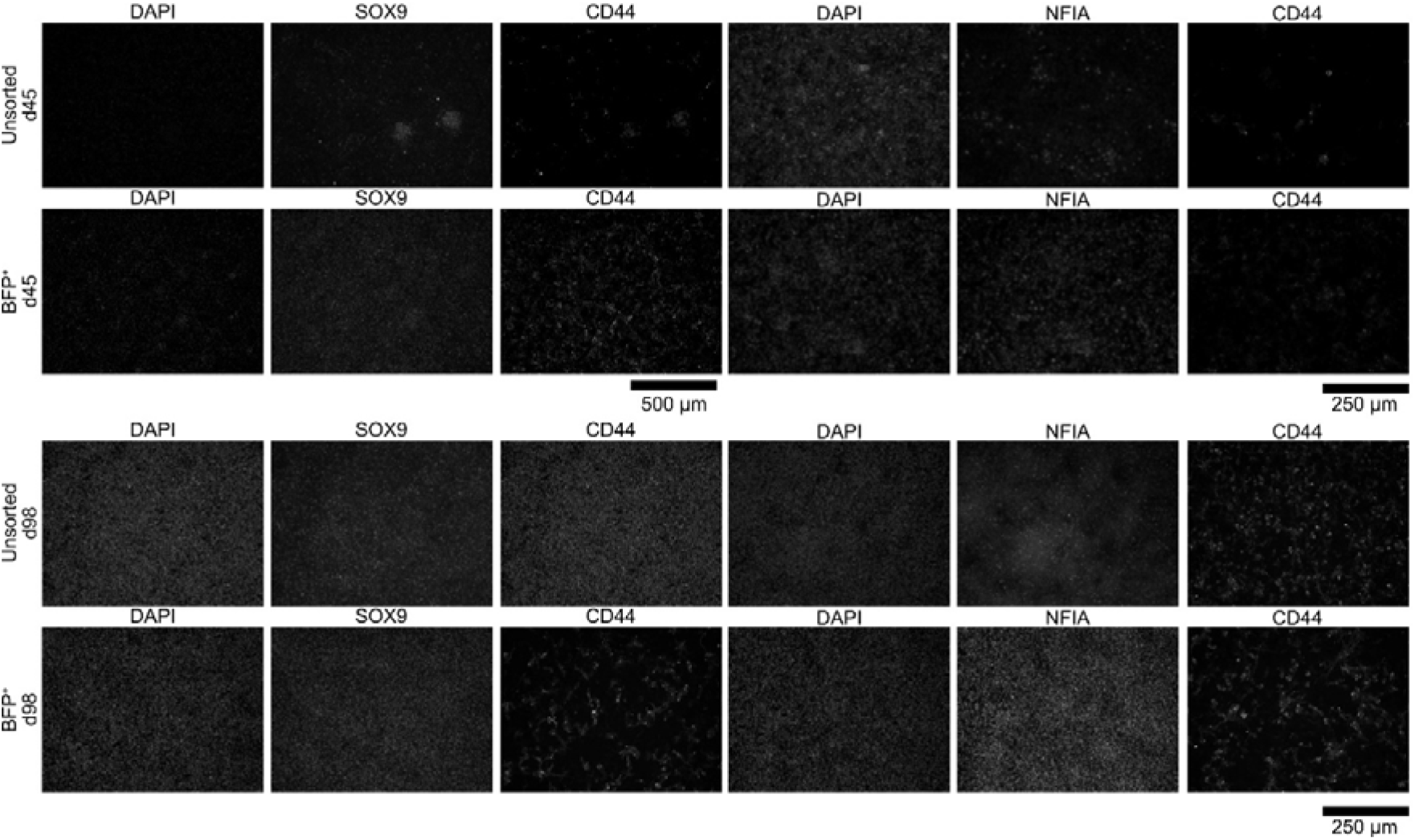
Original images of immunocytochemistry of astrogenic markers shown in Figure 2A.

**Figure S3.**
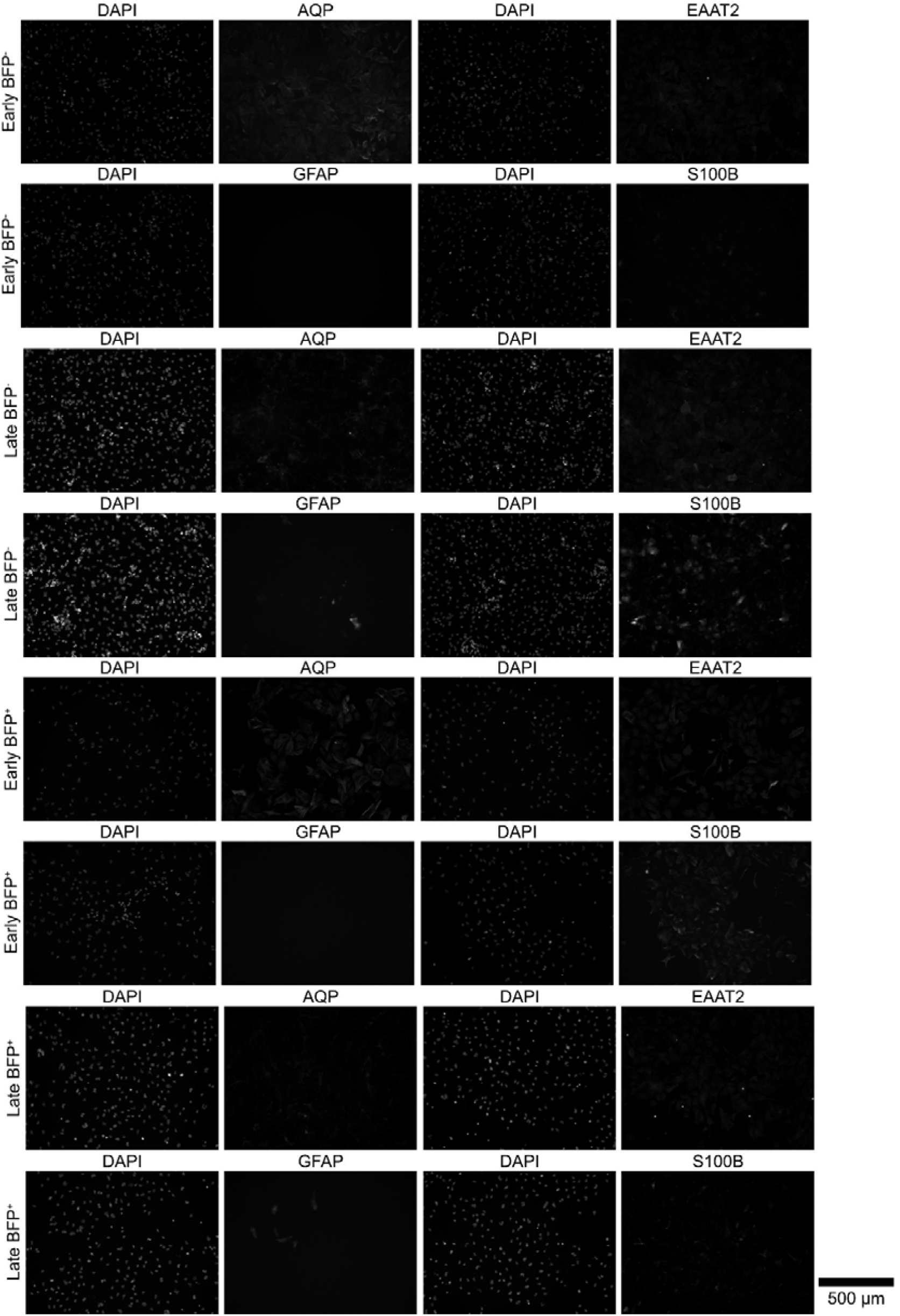
Original images of immunocytochemistry of astrocyte markers shown in Figure 2B.

**Figure S4.**
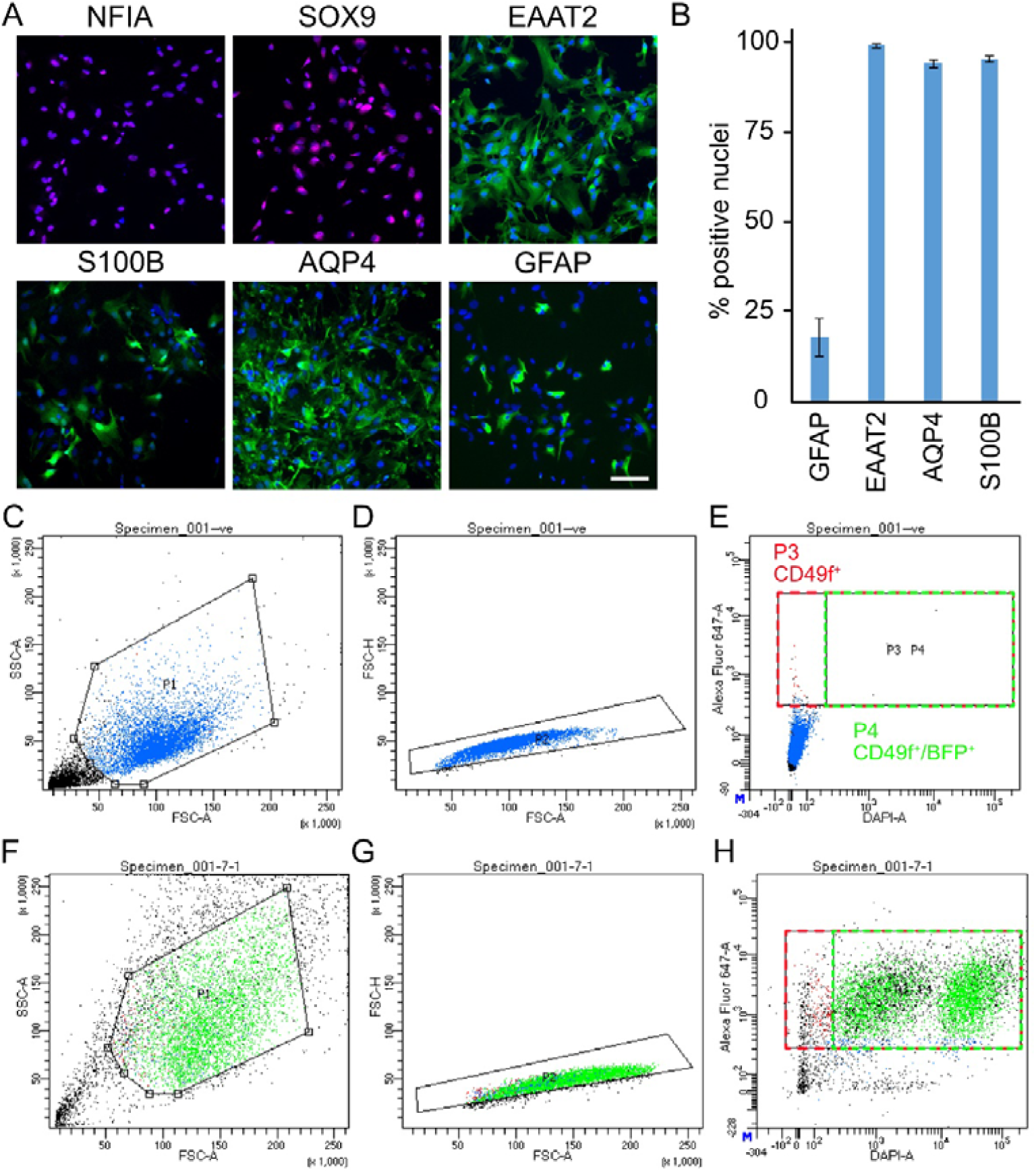
Non-patterned astrocytes and fluorescence-activated cell sorting of astrocytes for single cell RNA sequencing. A, Representative view of immunocytochemistry of astrocyte marker expression in non-patterned astrocytes (scale bar represents 100 µm). B, Quantification of astrocyte marker expression in astrocytes. Error bars represent SEM of three independent experiments. C-E, negative control samples; F-H, one sample of BFP+ astrocytes). C and F shows the dot plot of SSC-A versus FSC-A. D and G shows the dot plot of FSC-H versus FSC-A. E and H shows the dot plot of Alexa Fluor-647-A (labelling CD49f) versus DAPI-A (labelling BFP). P3 was used to isolate CD49f^+^ population (including both BFP^+^ and BFP^-^), while P4 was used to isolate CD49f^+^/BFP^+^ population.

**Figure S5.**
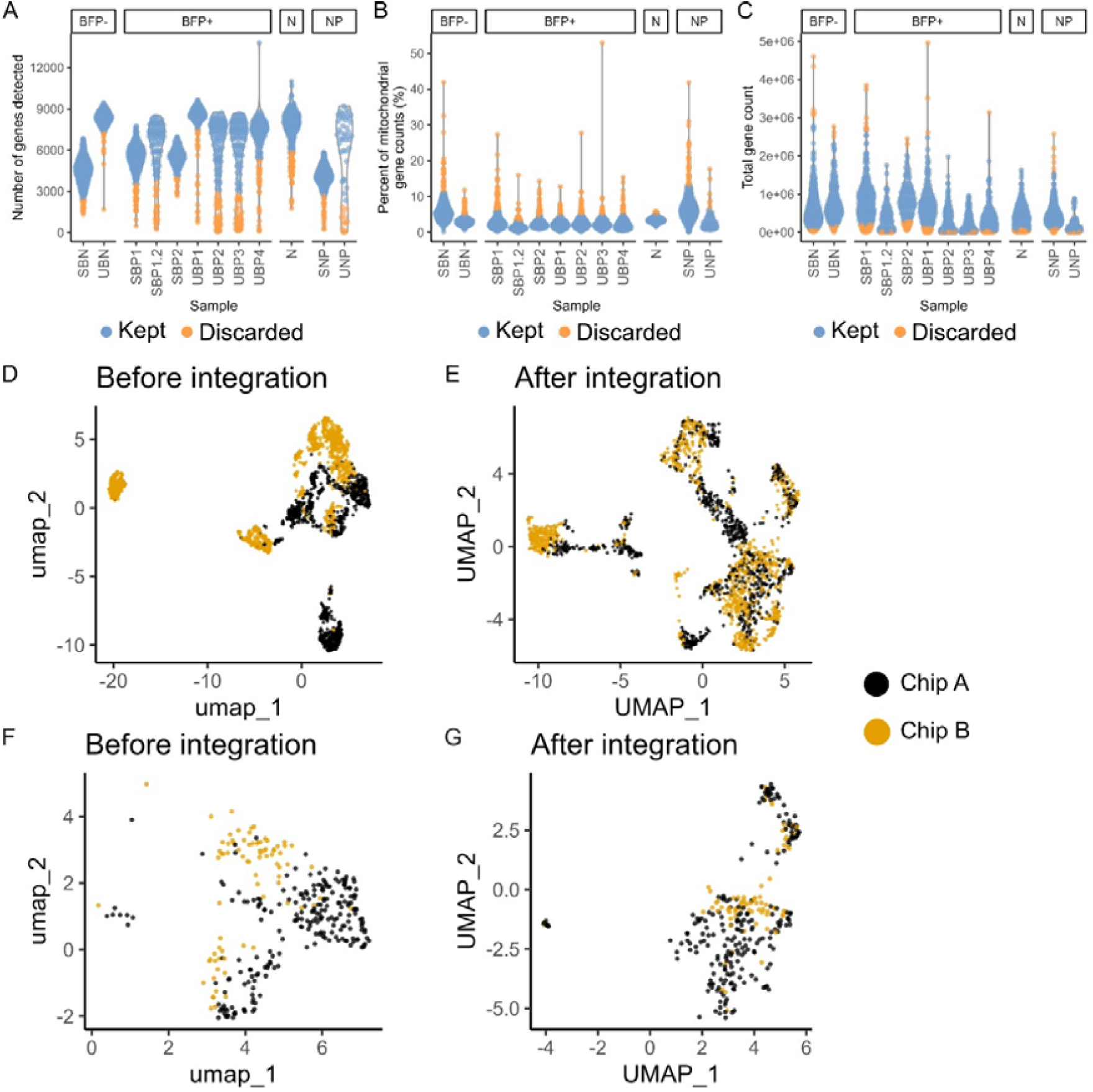
Processing of single cell RNA sequencing data. A-C , Violin plots showing the results of adaptive cell-level filtering based on the number of genes detected per cell (A), percentage of mitochondrial gene counts per cell (B), and total gene count per cell (C). D-E, UMAP plot of all filtered cells before (D) and after integration (E) coloured by chip. F-G, UMAP plot of the subset of sorted BFP^+^ astrocytes on Chip A and B derived from the same astrocyte differentiation before (F) and after integration (G) coloured by chip. (N: neuron; SBN: sorted BFP^-^; SBP: sorted BFP^+^; SNP: sorted non-patterned; UBN: unsorted BFP^-^; UBP: unsorted BFP^+^; UNP: unsorted non-patterned; numbers represent independent samples)

**Figure S6.**
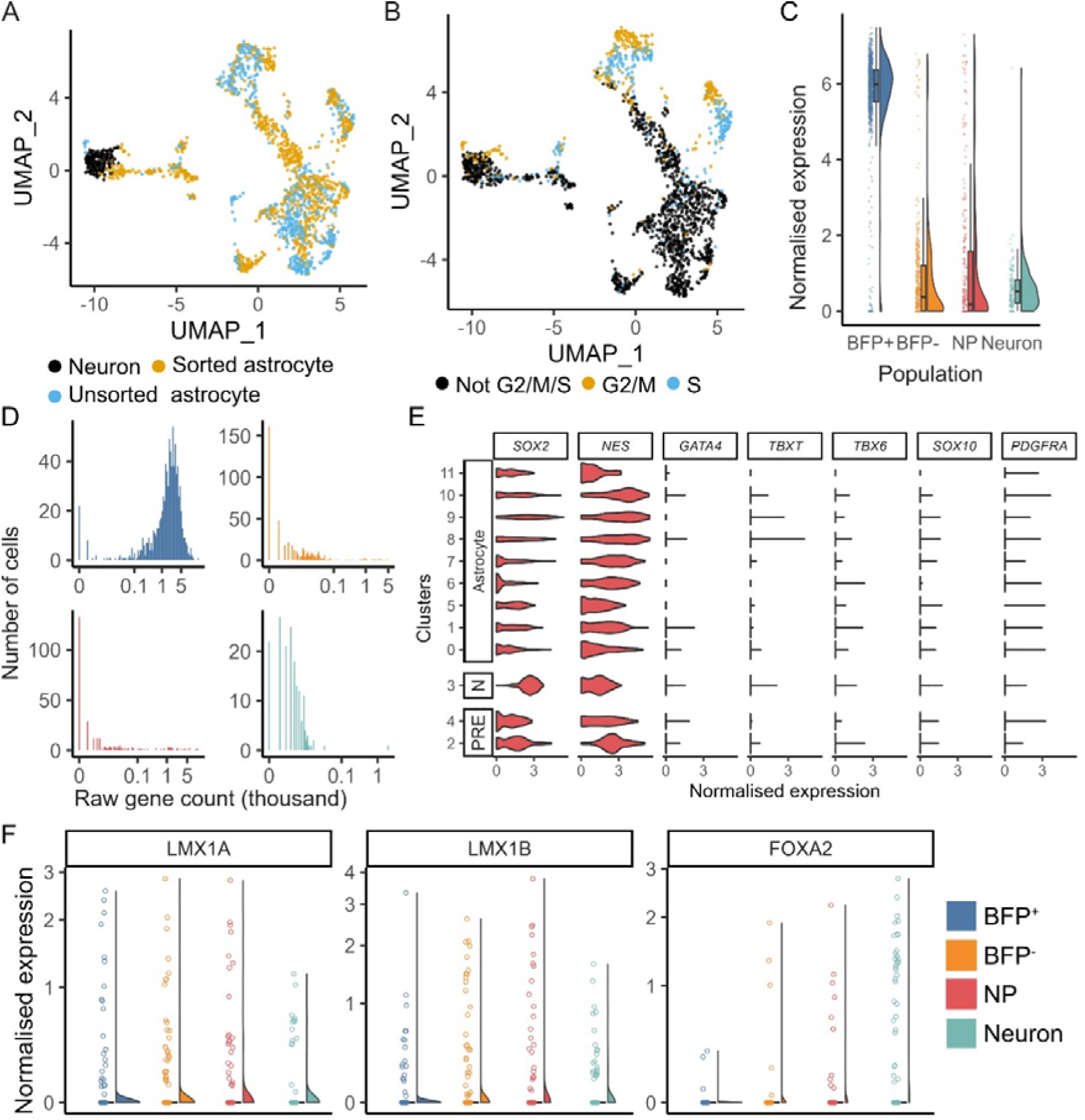
Marker expression in scRNAseq. A-B, UMAP plot of all filtered cells after integration coloured by sample type and sorting status (A) and estimated cell cycle phase (B). C, Raincloud plots of normalised expression of *BFP*. D, Histogram of the distribution of raw gene count of *BFP*. E, Violin plots of the normalised expression of endoderm, mesoderm, neuroectoderm, and oligodendrocyte progenitor markers. F, Violin plots of the normalised expression of classic ventral midbrain markers.

**Figure S7.**
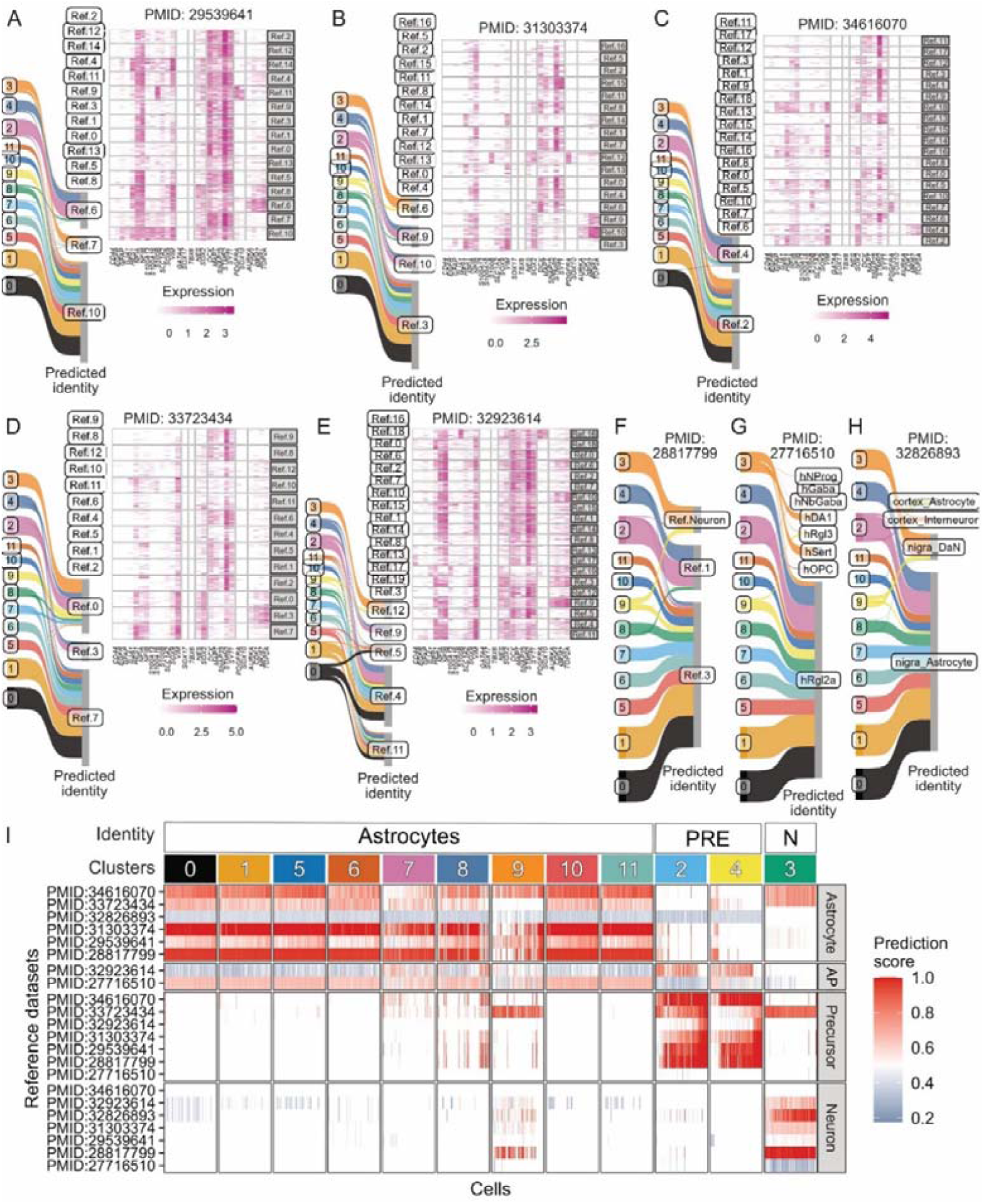
Reference mapping to human brain single cell RNA sequencing datasets. A-H , Sankey plot summarising the result of reference mapping of cells in different clusters to eight published reference human brain astrocyte scRNAseq datasets. The thickness of the thread is proportional to the number of cells mapped to the same identity in the reference datasets (predicted identity). Cluster IDs in this study are shown on the left. Heatmap in Panel A-E shows the expression of marker genes in different clusters in the re-annotated reference datasets. I, Heatmap of prediction score from Seurat integration.

**Figure S8.**
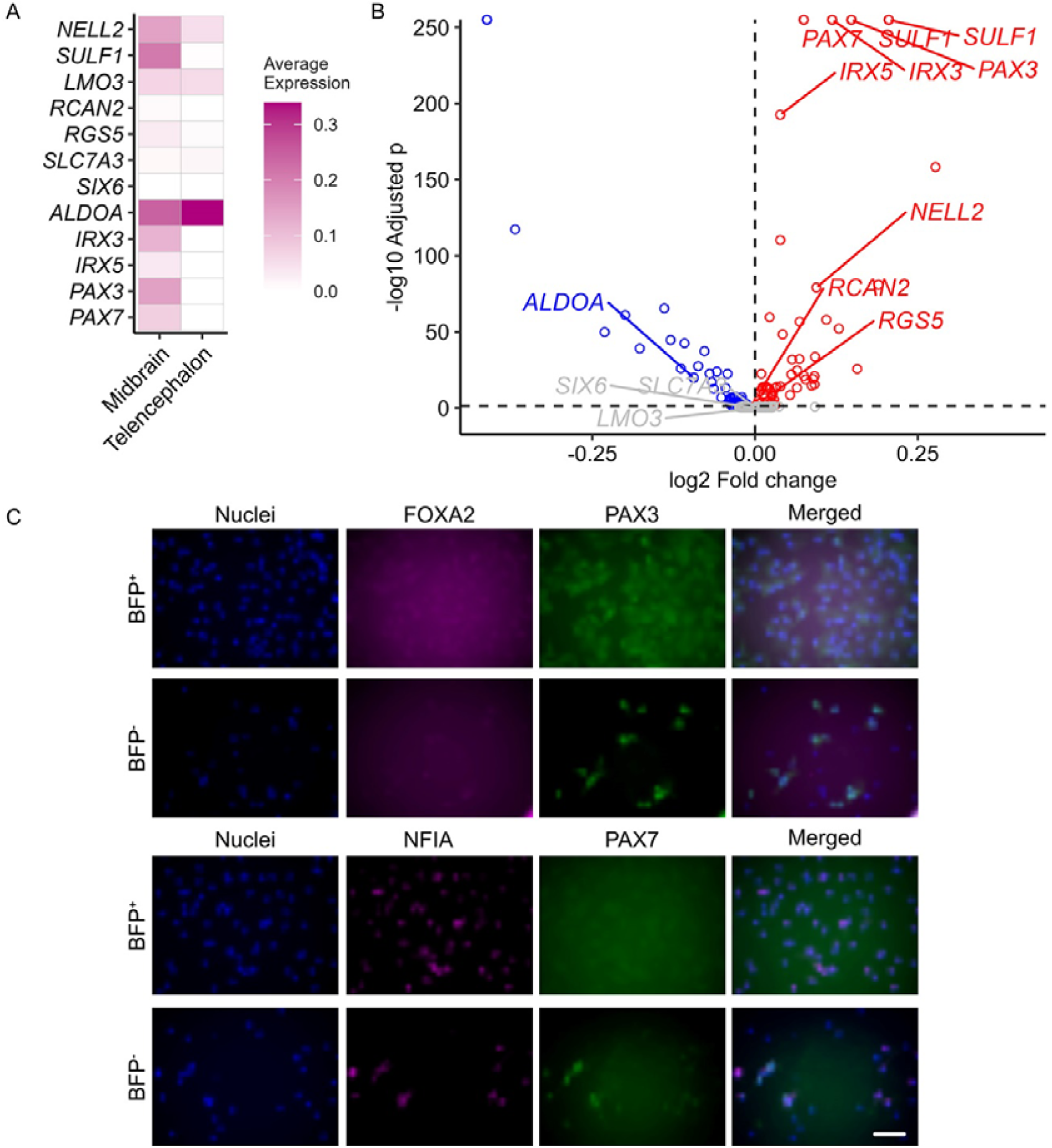
Validation of candidate markers for BFP+ and BFP-astrocytes. A, Heatmap of the average expression of candidate markers in mesencephalon and telencephalon astrocytes from five human foetal brain scRNAseq datasets. B, Volcano plot showing the log2 fold-change and adjusted p values of differential gene expression analysis comparing candidate markers in mesencephalon and telencephalon astrocytes from five human foetal brain scRNAseq datasets. Positive log2 fold-change represents higher expression level in mesencephalon astrocytes. C, Immunocytochemistry validation of PAX3, PAX7, and FOXA2 expression in BFP+ and BFP-astrocytes. (Scale bar represents 100 µm).

**Supplementary Table 1.**
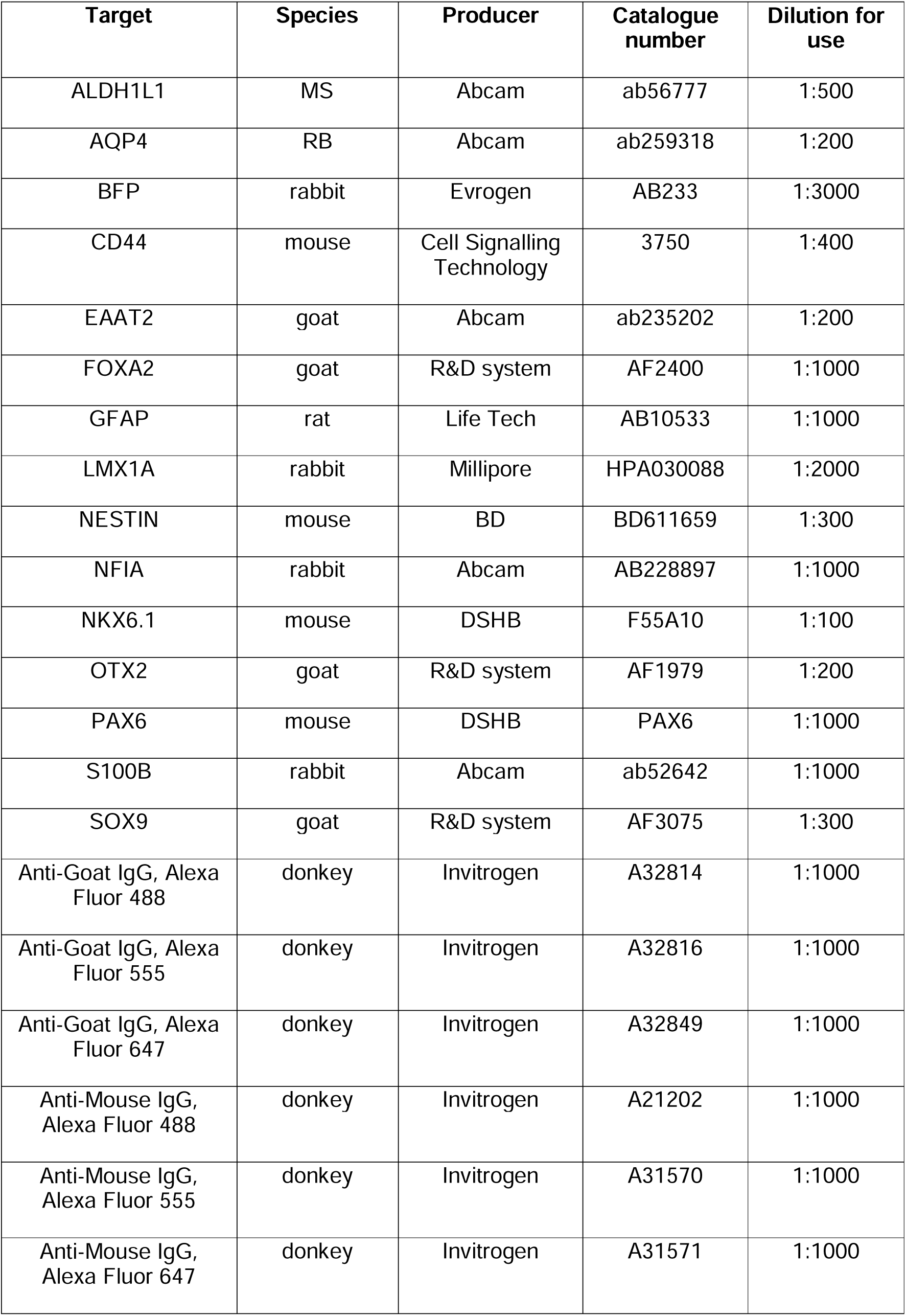
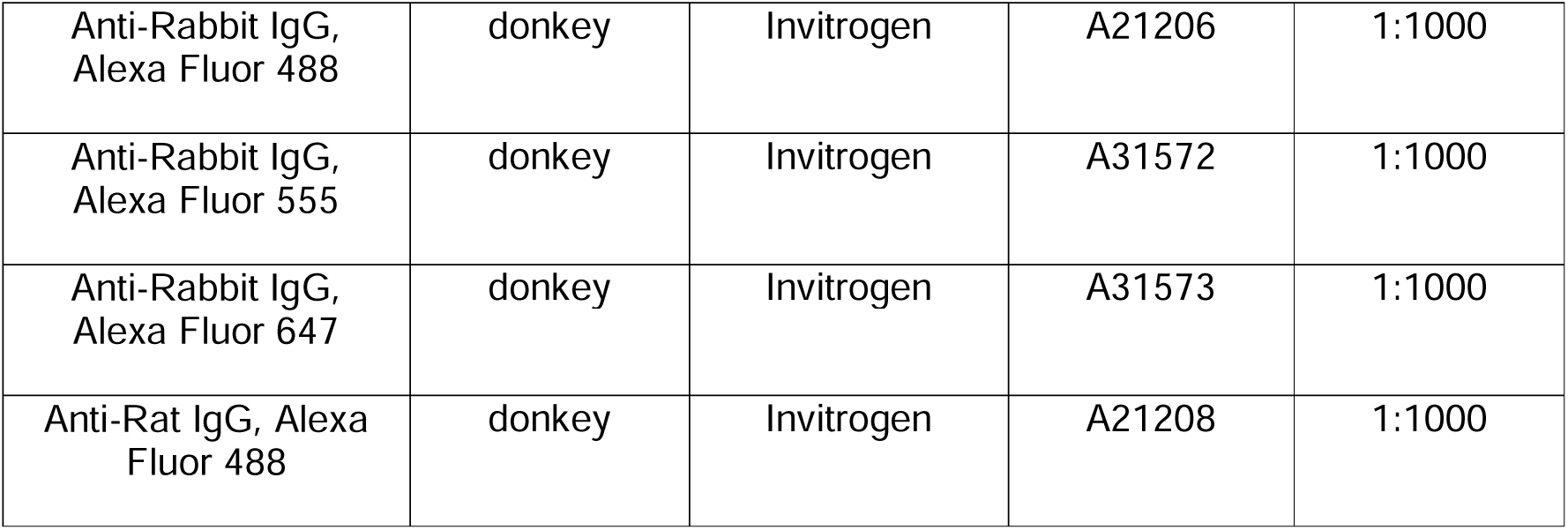
Antibodies used in this study.

**Supplementary Table 2.**
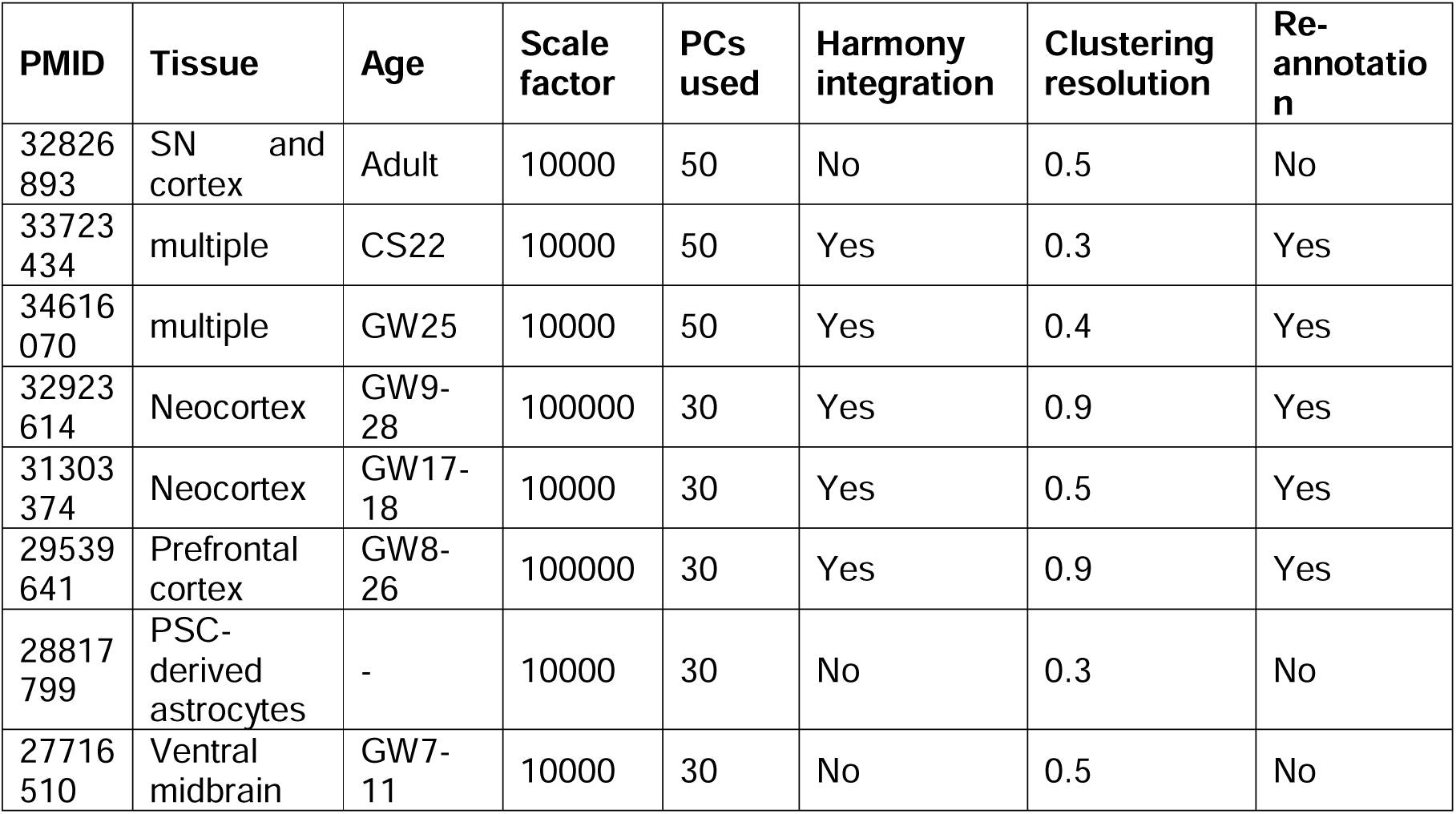
Settings used for processing published datasets.

Supplementary Data 1: Pairwise DEGs of individual astrocyte population with neurons.

Supplementary Data 2: GO enrichment of population enriched DEGs.

Supplementary Data 3: Representative enriched GO terms of DEGs for BFP+ and BFP-astrocytes.

Supplementary Data 4: DEGs comparing sorted and unsorted populations.

Supplementary Date 5: In silico validation of BFP^+^ and BFP^-^ astrocyte DEGs in integrated human foetal midbrain and forebrain scRNAseq datasets.

## References

Agarwal, D. et al. 2020. A single-cell atlas of the human substantia nigra reveals cell-specific pathways associated with neurological disorders. Nat Commun 11(1), p. 4183. doi: 10.1038/s41467-020-17876-0

Ahmed, M., Owens, M. J. S., Toledo, E. M., Arenas, E., Bradley, M. and Ffrench-Constant, C. 2021. Combinatorial ECM Arrays Identify Cooperative Roles for Matricellular Proteins in Enhancing the Generation of TH+ Neurons From Human Pluripotent Cells. Front Cell Dev Biol 9, p. 755406. doi: 10.3389/fcell.2021.755406

Andersson, E. et al. 2006. Identification of intrinsic determinants of midbrain dopamine neurons. Cell 124(2), pp. 393–405. doi: 10.1016/j.cell.2005.10.037

Ang, S. L., Wierda, A., Wong, D., Stevens, K. A., Cascio, S., Rossant, J. and Zaret, K. S. 1993. The formation and maintenance of the definitive endoderm lineage in the mouse: Involvement of HNF3/forkhead proteins. Development 119(4), pp. 1301–1315.

Barbar, L. et al. 2020. CD49f Is a Novel Marker of Functional and Reactive Human iPSC-Derived Astrocytes. Neuron 107(3), pp. 436–453 e412. doi: 10.1016/j.neuron.2020.05.014

Barbuti, P. A. et al. 2020. IPSC-derived midbrain astrocytes from Parkinson’s disease patients carrying pathogenic SNCA mutations exhibit alpha-synuclein aggregation, mitochondrial fragmentation and excess calcium release. bioRxiv, p. 2020.2004.2027.053470. doi: 10.1101/2020.04.27.053470

Batiuk, M. Y. et al. 2020. Identification of region-specific astrocyte subtypes at single cell resolution. Nature Communications 11(1), doi: 10.1038/s41467-019-14198-8

Bayraktar, O. A. et al. 2020. Astrocyte layers in the mammalian cerebral cortex revealed by a single-cell in situ transcriptomic map. Nature Neuroscience 23(4), pp. 500–509. doi: 10.1038/s41593-020-0602-1

Bhaduri, A. et al. 2021. An atlas of cortical arealization identifies dynamic molecular signatures. Nature 598(7879), pp. 200–204. doi: 10.1038/s41586-021-03910-8

Bifsha, P., Balsalobre, A. and Drouin, J. 2017. Specificity of Pitx3-Dependent Gene Regulatory Networks in Subsets of Midbrain Dopamine Neurons. Molecular Neurobiology 54(7), pp. 4921–4935. doi: 10.1007/s12035-016-0040-y

Booth, H. D. E., Hirst, W. D. and Wade-Martins, R. 2017. The Role of Astrocyte Dysfunction in Parkinson’s Disease Pathogenesis. Trends in neurosciences 40(6), pp. 358–370. doi: 10.1016/j.tins.2017.04.001

Booth, H. D. E. et al. 2019. RNA sequencing reveals MMP2 and TGFB1 downregulation in LRRK2 G2019S Parkinson’s iPSC-derived astrocytes. Neurobiol Dis 129, pp. 56–66. doi: 10.1016/j.nbd.2019.05.006

Bradley, R. A. et al. 2019. Regionally specified human pluripotent stem cell-derived astrocytes exhibit different molecular signatures and functional properties. Development 146(13), p. dev170910. doi: 10.1242/dev.170910

Cardo, L. F., Monzón-Sandoval, J., Li, Z., Webber, C. and Li, M. 2023. Single-Cell Transcriptomics and In Vitro Lineage Tracing Reveals Differential Susceptibility of Human iPSC-Derived Midbrain Dopaminergic Neurons in a Cellular Model of Parkinson’s Disease. Cells 12(24), doi: 10.3390/cells12242860

Carlson, M. 2019. org.Hs.eg.db: Genome wide annotation for Human. R package version 3.8.2.

Chai, H. et al. 2017. Neural Circuit-Specialized Astrocytes: Transcriptomic, Proteomic, Morphological, and Functional Evidence. Neuron 95(3), pp. 531–549 e539. doi: 10.1016/j.neuron.2017.06.029

Chandrasekaran, A., Avci, H. X., Leist, M., Kobolák, J. and Dinnyés, A. 2016. Astrocyte Differentiation of Human Pluripotent Stem Cells: New Tools for Neurological Disorder Research. Frontiers in cellular neuroscience 10, pp. 215–215. doi: 10.3389/fncel.2016.00215

Chung, N. C. and Storey, J. D. 2015. Statistical significance of variables driving systematic variation in high-dimensional data. Bioinformatics 31(4), pp. 545–554. doi: 10.1093/bioinformatics/btu674

Clough, E. and Barrett, T. 2016. The Gene Expression Omnibus Database. Methods Mol Biol 1418, pp. 93–110. doi: 10.1007/978-1-4939-3578-9_5

Crompton, L. A., McComish, S. F., Stathakos, P., Cordero-Llana, O., Lane, J. D. and Caldwell, M. A. 2021. Efficient and Scalable Generation of Human Ventral Midbrain Astrocytes from Human-Induced Pluripotent Stem Cells. Journal of Visualized Experiments (176), doi: 10.3791/62095

Crompton, L. A., McComish, S. F., Steward, T. G. J., Whitcomb, D. J., Lane, J. D. and Caldwell, M. A. 2023. Human stem cell-derived ventral midbrain astrocytes exhibit a region-specific secretory profile. Brain Commun 5(2), p. fcad114. doi: 10.1093/braincomms/fcad114

de Rus Jacquet, A., Tancredi, J. L., Lemire, A. L., DeSantis, M. C., Li, W. P. and O’Shea, E. K. 2021. The LRRK2 G2019S mutation alters astrocyte-to-neuron communication via extracellular vesicles and induces neuron atrophy in a human iPSC-derived model of Parkinson’s disease. Elife 10, p. 2020.2007.2002.178574. doi: 10.7554/eLife.73062

Deneen, B., Ho, R., Lukaszewicz, A., Hochstim, C. J., Gronostajski, R. M. and Anderson, D. J. 2006. The Transcription Factor NFIA Controls the Onset of Gliogenesis in the Developing Spinal Cord. Neuron 52(6), pp. 953–968. doi: 10.1016/j.neuron.2006.11.019

di Domenico, A. et al. 2019. Patient-Specific iPSC-Derived Astrocytes Contribute to Non-Cell-Autonomous Neurodegeneration in Parkinson’s Disease. Stem cell reports 12(2), pp. 213–229. doi: 10.1016/j.stemcr.2018.12.011

Dobin, A. et al. 2013. STAR: ultrafast universal RNA-seq aligner. Bioinformatics 29(1), pp. 15–21. doi: 10.1093/bioinformatics/bts635

Duan, D., Fu, Y., Paxinos, G. and Watson, C. 2013. Spatiotemporal expression patterns of Pax6 in the brain of embryonic, newborn, and adult mice. Brain Structure and Function 218(2), pp. 353–372. doi: 10.1007/s00429-012-0397-2

Endo, F. et al. 2022. Molecular basis of astrocyte diversity and morphology across the CNS in health and disease. Science 378(6619), p. eadc9020. doi: doi:10.1126/science.adc9020

Eze, U. C., Bhaduri, A., Haeussler, M., Nowakowski, T. J. and Kriegstein, A. R. 2021. Single-cell atlas of early human brain development highlights heterogeneity of human neuroepithelial cells and early radial glia. Nature Neuroscience 24(4), pp. 584–594. doi: 10.1038/s41593-020-00794-1

Failli, V., Bachy, I. and Rétaux, S. 2002. Expression of the LIM-homeodomain gene Lmx1a (dreher) during development of the mouse nervous system. Mechanisms of Development 118(1-2), pp. 225–228. doi: 10.1016/s0925-4773(02)00254-x

Falk, A. et al. 2012. Capture of neuroepithelial-like stem cells from pluripotent stem cells provides a versatile system for in vitro production of human neurons. PLOS ONE 7(1), p. e29597. doi: 10.1371/journal.pone.0029597

Fan, X. et al. 2020. Single-cell transcriptome analysis reveals cell lineage specification in temporal-spatial patterns in human cortical development. Science Advances 6(34), p. eaaz2978. doi: doi:10.1126/sciadv.aaz2978

Finak, G. et al. 2015. MAST: a flexible statistical framework for assessing transcriptional changes and characterizing heterogeneity in single-cell RNA sequencing data. Genome Biology 16(1), p. 278. doi: 10.1186/s13059-015-0844-5

Gabay, L., Lowell, S., Rubin, L. L. and Anderson, D. J. 2003. Deregulation of dorsoventral patterning by FGF confers trilineage differentiation capacity on CNS stem cells in vitro. Neuron 40(3), pp. 485–499. doi: 10.1016/s0896-6273(03)00637-8

Hedegaard, A., Monzón-Sandoval, J., Newey, S. E., Whiteley, E. S., Webber, C. and Akerman, C. J. 2020. Pro-maturational Effects of Human iPSC-Derived Cortical Astrocytes upon iPSC-Derived Cortical Neurons. Stem cell reports 15(1), pp. 38–51. doi: 10.1016/j.stemcr.2020.05.003

Holmqvist, S. et al. 2015. Generation of human pluripotent stem cell reporter lines for the isolation of and reporting on astrocytes generated from ventral midbrain and ventral spinal cord neural progenitors. Stem Cell Research 15(1), pp. 203–220. doi: 10.1016/j.scr.2015.05.014

Houweling, A. C., Dildrop, R., Peters, T., Mummenhoff, J., Moorman, A. F. M., Rüther, U. and Christoffels, V. M. 2001. Gene and cluster-specific expression of the Iroquois family members during mouse development. Mechanisms of Development 107(1), pp. 169–174. doi: 10.1016/S0925-4773(01)00451-8

Itoh, N. et al. 2018. Cell-specific and region-specific transcriptomics in the multiple sclerosis model: Focus on astrocytes. Proceedings of the National Academy of Sciences 115(2), pp. E302–E309. doi: doi:10.1073/pnas.1716032115

Jaeger, I. et al. 2011. Temporally controlled modulation of FGF/ERK signaling directs midbrain dopaminergic neural progenitor fate in mouse and human pluripotent stem cells. Development 138(20), pp. 4363–4374. doi: 10.1242/dev.066746

Jain, M., Armstrong, R. J., Tyers, P., Barker, R. A. and Rosser, A. E. 2003. GABAergic immunoreactivity is predominant in neurons derived from expanded human neural precursor cells in vitro. Exp Neurol 182(1), pp. 113–123. doi: 10.1016/s0014-4886(03)00055-4

Kamath, T. et al. 2022. Single-cell genomic profiling of human dopamine neurons identifies a population that selectively degenerates in Parkinson’s disease. Nature Neuroscience 25(5), pp. 588–595. doi: 10.1038/s41593-022-01061-1

Koch, P., Opitz, T., Steinbeck, J. A., Ladewig, J. and Brüstle, O. 2009. A rosette-type, self-renewing human ES cell-derived neural stem cell with potential for in vitro instruction and synaptic integration. Proc Natl Acad Sci U S A 106(9), pp. 3225–3230. doi: 10.1073/pnas.0808387106

Kostuk, E. W., Cai, J. and Iacovitti, L. 2019. Subregional differences in astrocytes underlie selective neurodegeneration or protection in Parkinson’s disease models in culture. Glia 67(8), pp. 1542–1557. doi: 10.1002/glia.23627

Krencik, R., Weick, J. P., Liu, Y., Zhang, Z.-J. and Zhang, S.-C. 2011. Specification of transplantable astroglial subtypes from human pluripotent stem cells. Nature Biotechnology 29, p. 528. doi: 10.1038/nbt.1877 https://www.nature.com/articles/nbt.1877#supplementary-information

La Manno, G. et al. 2016. Molecular Diversity of Midbrain Development in Mouse, Human, and Stem Cells. Cell 167(2), pp. 566–580.e519. doi: 10.1016/j.cell.2016.09.027

Li, Y. et al. 2023. Spatiotemporal transcriptome atlas reveals the regional specification of the developing human brain. Cell 186(26), pp. 5892–5909.e5822. doi: 10.1016/j.cell.2023.11.016

Liao, Y., Smyth, G. K. and Shi, W. 2013. The Subread aligner: fast, accurate and scalable read mapping by seed-and-vote. Nucleic Acids Res 41(10), p. e108. doi: 10.1093/nar/gkt214

Lin, Y.-T. et al. 2018. APOE4 Causes Widespread Molecular and Cellular Alterations Associated with Alzheimer’s Disease Phenotypes in Human iPSC-Derived Brain Cell Types. Neuron 98(6), pp. 1141–1154.e1147. doi: 10.1016/j.neuron.2018.05.008

Liu, Y. et al. 2004. CD44 expression identifies astrocyte-restricted precursor cells. Developmental Biology 276(1), pp. 31–46. doi: 10.1016/j.ydbio.2004.08.018

Lozzi, B., Huang, T. W., Sardar, D., Huang, A. Y. S. and Deneen, B. 2020. Regionally Distinct Astrocytes Display Unique Transcription Factor Profiles in the Adult Brain. Frontiers in Neuroscience 14, doi: 10.3389/fnins.2020.00061

Makarava, N., Chang, J. C.-Y., Kushwaha, R. and Baskakov, I. V. 2019. Region-Specific Response of Astrocytes to Prion Infection. Frontiers in Neuroscience 13, doi: 10.3389/fnins.2019.01048

Matsunaga, E., Araki, I. and Nakamura, H. 2001. Role of Pax3/7 in the tectum regionalization. Development 128(20), pp. 4069–4077. doi: 10.1242/dev.128.20.4069

McCarthy, D. J., Campbell, K. R., Lun, A. T. L. and Wills, Q. F. 2017. Scater: pre-processing, quality control, normalization and visualization of single-cell RNA-seq data in R. Bioinformatics 33(8), pp. 1179–1186. doi: 10.1093/bioinformatics/btw777

Molofsky, A. V. et al. 2012. Astrocytes and disease: a neurodevelopmental perspective. Genes & Development 26(9), pp. 891–907. doi: 10.1101/gad.188326.112

Morel, L. et al. 2017. Molecular and Functional Properties of Regional Astrocytes in the Adult Brain. J Neurosci 37(36), pp. 8706–8717. doi: 10.1523/JNEUROSCI.3956-16.2017

Nolbrant, S., Heuer, A., Parmar, M. and Kirkeby, A. 2017. Generation of high-purity human ventral midbrain dopaminergic progenitors for in vitro maturation and intracerebral transplantation. Nature Protocols 12(9), pp. 1962–1979. doi: 10.1038/nprot.2017.078

Oberheim, N. A. et al. 2009. Uniquely hominid features of adult human astrocytes. J Neurosci 29(10), pp. 3276–3287. doi: 10.1523/jneurosci.4707-08.2009

Peteri, U.-K., Pitkonen, J., Utami, K. H., Paavola, J., Roybon, L., Pouladi, M. A. and Castrén, M. L. 2021. Generation of the Human Pluripotent Stem-Cell-Derived Astrocyte Model with Forebrain Identity. Brain sciences 11(2), p. 209. doi: 10.3390/brainsci11020209

Phatnani, H. and Maniatis, T. 2015. Astrocytes in neurodegenerative disease. Cold Spring Harbor perspectives in biology 7(6), p. a020628. doi: 10.1101/cshperspect.a020628

Polioudakis, D. et al. 2019. A Single-Cell Transcriptomic Atlas of Human Neocortical Development during Mid-gestation. Neuron 103(5), pp. 785–801.e788. doi: 10.1016/j.neuron.2019.06.011

Roybon, L. et al. 2013. Human stem cell-derived spinal cord astrocytes with defined mature or reactive phenotypes. Cell Rep 4(5), pp. 1035–1048. doi: 10.1016/j.celrep.2013.06.021

Schindelin, J. et al. 2012. Fiji: an open-source platform for biological-image analysis. Nature Methods 9(7), pp. 676–682. doi: 10.1038/nmeth.2019

Schlicker, A., Domingues, F. S., Rahnenführer, J. and Lengauer, T. 2006. A new measure for functional similarity of gene products based on Gene Ontology. BMC Bioinformatics 7(1), p. 302. doi: 10.1186/1471-2105-7-302

Schober, A. L., Wicki-Stordeur, L. E., Murai, K. K. and Swayne, L. A. 2022. Foundations and implications of astrocyte heterogeneity during brain development and disease. Trends in neurosciences 45(9), pp. 692–703. doi: 10.1016/j.tins.2022.06.009

Serio, A. et al. 2013. Astrocyte pathology and the absence of non-cell autonomy in an induced pluripotent stem cell model of TDP-43 proteinopathy. Proceedings of the National Academy of Sciences of the United States of America 110(12), pp. 4697–4702. doi: 10.1073/pnas.1300398110

Sloan, S. A. et al. 2017. Human Astrocyte Maturation Captured in 3D Cerebral Cortical Spheroids Derived from Pluripotent Stem Cells. Neuron 95(4), pp. 779–790 e776. doi: 10.1016/j.neuron.2017.07.035

Sonninen, T.-M. et al. 2020. Metabolic alterations in Parkinson’s disease astrocytes. Scientific Reports 10(1), p. 14474. doi: 10.1038/s41598-020-71329-8

Stirling, D. R., Swain-Bowden, M. J., Lucas, A. M., Carpenter, A. E., Cimini, B. A. and Goodman, A. 2021. CellProfiler 4: improvements in speed, utility and usability. BMC Bioinformatics 22(1), p. 433. doi: 10.1186/s12859-021-04344-9

Stolt, C. C., Lommes, P., Sock, E., Chaboissier, M.-C., Schedl, A. and Wegner, M. 2003. The Sox9 transcription factor determines glial fate choice in the developing spinal cord. Genes & Development 17(13), pp. 1677–1689. doi: 10.1101/gad.259003

Strelau, J. et al. 2000. Growth/differentiation factor-15/macrophage inhibitory cytokine-1 is a novel trophic factor for midbrain dopaminergic neurons in vivo. J Neurosci 20(23), pp. 8597–8603. doi: 10.1523/jneurosci.20-23-08597.2000

Stuart, T. et al. 2019. Comprehensive Integration of Single-Cell Data. Cell 177(7), pp. 1888–1902.e1821. doi: 10.1016/j.cell.2019.05.031

Sun, Y. et al. 2008. Long-term tripotent differentiation capacity of human neural stem (NS) cells in adherent culture. Mol Cell Neurosci 38(2), pp. 245–258. doi: 10.1016/j.mcn.2008.02.014

Supek, F., Bošnjak, M., Škunca, N. and Šmuc, T. 2011. REVIGO Summarizes and Visualizes Long Lists of Gene Ontology Terms. PLOS ONE 6(7), p. e21800. doi: 10.1371/journal.pone.0021800

Takata, N. and Hirase, H. 2008. Cortical Layer 1 and Layer 2/3 Astrocytes Exhibit Distinct Calcium Dynamics In Vivo. PLOS ONE 3(6), p. e2525. doi: 10.1371/journal.pone.0002525

Tcw, J. et al. 2017. An Efficient Platform for Astrocyte Differentiation from Human Induced Pluripotent Stem Cells. Stem cell reports 9(2), pp. 600–614. doi: 10.1016/j.stemcr.2017.06.018

Team, R. C. 2023. : A Language and Environment for Statistical Computing. Vienna, Austria: R Foundation for Statistical Computing, .

Verkhratsky, A. and Nedergaard, M. 2018. Physiology of astroglia. Physiological Reviews 98(1), pp. 239–389. doi: 10.1152/physrev.00042.2016

Xin, W., Schuebel, K. E., Jair, K.-w., Cimbro, R., De Biase, L. M., Goldman, D. and Bonci, A. 2019. Ventral midbrain astrocytes display unique physiological features and sensitivity to dopamine D2 receptor signaling. Neuropsychopharmacology 44(2), pp. 344–355. doi: 10.1038/s41386-018-0151-4

Yang, Y. et al. 2022. Therapeutic functions of astrocytes to treat α-synuclein pathology in Parkinson’s disease. Proceedings of the National Academy of Sciences 119(29), p. e2110746119. doi: doi:10.1073/pnas.2110746119

Yun, W. et al. 2019. Generation of Anterior Hindbrain-Specific, Glial-Restricted Progenitor-Like Cells from Human Pluripotent Stem Cells. Stem Cells and Development 28(10), pp. 633–648. doi: 10.1089/scd.2019.0033

Zhong, S. et al. 2018. A single-cell RNA-seq survey of the developmental landscape of the human prefrontal cortex. Nature 555(7697), pp. 524–528. doi: 10.1038/nature25980

Zhou, S. et al. 2016. Neurosphere Based Differentiation of Human iPSC Improves Astrocyte Differentiation. Stem Cells Int 2016, p. 4937689. doi: 10.1155/2016/4937689

Zhu, Q., Shah, S., Dries, R., Cai, L. and Yuan, G. C. 2018. Identification of spatially associated subpopulations by combining scRNAseq and sequential fluorescence in situ hybridization data. Nat Biotechnol 36(12), pp. 1183–1190. doi: 10.1038/nbt.4260

